# Subclinical anxiety is associated with reduced self-distancing and enhanced self-blame-related connectivity between anterior temporal and subgenual cingulate cortices

**DOI:** 10.1101/2025.09.20.677492

**Authors:** Michal Rafal Zareba, Ivan González-García, Marcos Ibáñez Montolio, Richard J. Binney, Paul Hoffman, Maya Visser

## Abstract

Excessive self-blaming emotions are commonly observed in anxiety disorders, with qualitatively similar symptomatology reported in subclinical populations. Interpretation of moral information requires assessing the social conceptual information, a process overseen by the superior anterior temporal lobe (sATL). Feelings of self-blame evoke interactions of sATL and socio-affective regions, and previous research shows that subclinical anxiety modulates the organisation of the self-blame circuitry. This study aimed to extend these findings by exploring links of trait-anxiety with (i) self-blaming emotions and associated behaviours in an experimental task, and (ii) self-blame-dependent neural activity and connectivity, as observed during reliving of autobiographical guilt memories. We also explored the role of resting-state fMRI in linking these phenomena. Increased anxiety was linked to stronger self-blaming emotions, and more pronounced self-attacking and hiding. When experiencing negative emotions about themselves (i.e. shame and self-anger), anxious individuals were also less likely to disengage from self-focused thoughts. These behavioural findings were paralleled by enhanced self-blame-related connectivity between the left sATL and bilateral posterior subgenual cingulate cortex. Distinct patterns of activity and connectivity within the ATL-related circuitry were furthermore linked to individual differences in intensity of the self-blaming emotions and approach-avoidance motivation towards the guilt memories. As such, the results of the current study link stronger self-blaming emotions in anxious individuals with specific maladaptive patterns of behaviour. Furthermore, the work provides robust evidence for the important role of ATL-related circuitry in self-blame processing, supporting its broader involvement in social conceptual processing and its alterations in subclinical anxiety.

## 1. Introduction

The growing prevalence of mood and anxiety disorders worldwide, a crisis that has accelerated in the years following the COVID-19 pandemic (Yuan et al., 2022), is a major challenge for modern society, as it comes with a high economic burden. Mental health problems significantly impact individual quality of life, with social difficulties being amongst the most often reported complaints. As navigating the complex world of interpersonal interactions is a key element of our everyday functioning, understanding the processes contributing to the emergence of these hardships is crucial. On one hand, it can inform novel, evidence-based strategies for improving social functioning of the affected individuals. On the other hand, it might contribute to the development of strategies preventing the onset of the problems in the first place.

Self-blaming emotions, such as guilt and shame, often arise in response to actual or predicted social punishment cues (Yang et al., 2024). Although they constitute a well established characteristic of mood and anxiety disorders (Beck, 1963; Cândea and Szentagotai-Tătar, 2018), they only recently gained attention from the neuroscientific community. The adaptive or maladaptive character of self-blaming emotions largely depends on the individual coping mechanisms (Janoff-Bulman, 1979; Tangney et al., 2007). They are considered adaptive when they lead to efforts aimed at making up for the negative consequences, as well as avoiding them in the future. In turn, the maladaptive forms of self-blaming are typically associated with social distancing and focus on being responsible for the outcomes, resulting in self-criticism and generalisation of such experiences to the self. Unsurprisingly, the prevalence of self-blaming emotions in major depressive disorder (MDD) patients has been previously linked to the maladaptive behavioural profile (Duan et al., 2021; Duan et al., 2023).

As for the neural basis of these observations, a number of neuroimaging studies highlighted the role of the right superior anterior temporal lobe regions involved in social semantics (sATL; Zahn et al., 2007; Thye et al., 2024). Specifically, reduced connectivity of the sATL with dorsolateral prefrontal cortex and ventral striatum during self-blaming was associated with the overgeneralisation of self-blaming emotions in in MDD patients (Green et al., 2013), while its coupling with subgenual anterior cingulate cortex (sgACC) was consistently found to be predictive of treatment response (Lythe et al., 2015; Zahn et al., 2019; Jaeckle et at., 2023; Fennema et al., 2023). Importantly, the connectivity with anterior and posterior parts of the sgACC was found to be differentially associated with depression progression. Given their functional differences, with the former linked to social agency and the latter to affiliative processing (Zahn, 2025), studying their coupling separately might provide insights into alterations in distinct self-blame-related processes.

Despite the fact that anxious individuals, similarly to MDD patients, display proneness to self-blaming emotions (Cândea and Szentagotai-Tătar, 2018), the behavioural and neural patterns associated with self-blaming in this group remain largely unexplored (González-García and Visser, 2023). In recent years it has been demonstrated that both subclinical and clinical anxious populations exhibit a comparable set of symptoms and similar underlying changes in neural activity (Besteher et al. 2017). Such accounts suggest that biologically-informed symptom-based models of mental health disorders may be more useful clinically than the current categorical classifications based on DSM-5-TR (Kas et al., 2025). For example, we demonstrated that increased generalised self-blaming of subclinically anxious individuals was primarily related to increased functional contributions of the left sATL to the guilt processing circuitry (Zareba et al, 2024). Such behaviours might be further reinforced by the disconnection of the inferior frontal and insular regions involved in cognitive emotion regulation, and a network structural architecture favouring local information processing (Zareba et al., 2024). Nevertheless, it still remains unknown whether the stronger self-blaming tendency observed in subclinically anxious individuals is associated with similar maladaptive behaviours and alterations in neural correlates of self-blaming experiences as previously reported in MDD patients.

Therefore, we set out to thoroughly investigate the self-blaming-related phenomena in subclinically anxious individuals, combining behavioural paradigms with functional magnetic resonance imaging (fMRI). We used the Moral Sentiment and Action Tendencies task (MSAT; Zahn et al., 2015; Duan et al., 2023) to study blame-related social feelings and associated action tendencies, and their dependency on whether one is the performer or a recipient of hypothetical inappropriate social actions (i.e. self- and other-agency). We expected to replicate prior findings in anxious individuals and observe higher levels of negative social feelings towards oneself (e.g., guilt and shame; Cândea and Szentagotai-Tătar, 2018). As the sense of agency plays a key role in the emergence of self-blaming emotions (Caspar et al., 2020), we additionally expected this phenomenon to be more pronounced on the self-agency trials. Regarding the action tendencies, we hypothesised that higher levels of anxiety would be associated with maladaptive behaviours, such as increased hiding (social avoidance) and self-attacking, similarly to MDD patients (Duan et al., 2021; Duan et al., 2023). We further presumed that these observations would be particularly prominent on the trials where one reports feeling guilt or shame (Tangney et al., 2007).

In addition, a subset of participants underwent fMRI while recalling autobiographical guilt memories, i.e. instances of strong self-blaming emotions. Apart from obtaining indices of neural activity and connectivity during self-blaming, we measured approach-avoidance and salience pertaining to the memories. As the MSAT task tests self-blaming emotions and associated action tendencies using hypothetical social scenarios, introduction of these two concepts in the guilt recollection task enabled measuring similar phenomena using highly emotional autobiographical events. Akin to the hypotheses for the MSAT task, we expected subclinically anxious individuals to report more avoidance towards their guilt memories, and perceive these events as more salient. On the neural level, we hypothesised that these associations would be paralleled by increased self-blame-dependent coupling between the sATL, which encodes meaning of social stimuli (Zahn et al., 2007; Binney et al., 2016), and posterior sgACC, which is involved in social affiliative processing (Zahn, 2025), mirroring the previous results in MDD patients (Lythe et al., 2015; Jaeckle et al., 2023).

Last but not least, building on our work using graph theory (Zareba et al., 2024), we aimed to characterise in greater detail the neural circuitry associated with self-blaming emotions, this time using complementary voxel-wise methodology. We obtained resting-state fMRI data, calculated measures of brain activity and connectivity, and finally correlated them with the strength of individual self-blaming emotions from the MSAT task (Zahn et al., 2015; Duan et al., 2023). We expected to observe statistically significant associations within the same set of regions as activated by the guilt recollection task deployed in this study, and particularly in the ATL (Zareba et al., 2024).

## 2. Methods

The current study included a behavioural task, fMRI with a task on autobiographical guilt memory processing, and resting-state fMRI. It adhered to the ethical standards of the Declaration of Helsinki and was approved by the local ethics committee (CEISH/07/2022). Participants provided informed consent prior to participation and were remunerated at a rate of 10 euros per hour.

### 2.1. Moral Sentiment and Action Tendencies Task (MSAT)

#### 2.1.1. Participants

The MSAT task was performed by 140 healthy volunteers recruited mainly from the university community via local announcements and word of mouth. As the participants were also meant to subsequently undergo magnetic resonance imaging (MRI), the typical inclusion criteria for MRI studies were applied: right-handedness as assessed by the Edinburgh Handedness Inventory (Oldfield, 1971), normal or corrected to normal vision, no history of neurological and psychiatric disorders, and not taking any psychotropic medication. Three measures of anxiety and associated facets were applied: State-Trait Anxiety Inventory (STAI; Spielberger, 1989), Sensitivity to Punishment and Sensitivity to Reward Questionnaire (SCSRQ; Torrubia et al., 2001) and Behavioral Inhibition and Behavioral Activation System (BIS/BAS; Carver and White, 1994), following the recommendation to assess symptomatology across several scales to increase results reliability and replicability (Fried and Nesse, 2015). For each participant, we calculated a composite score of anxiety by standardising (Z-score) their scores for trait-anxiety, sensitivity to punishment and behavioural inhibition system, and adding them all together, following the methodology established in previous studies (Holmes et al., 2012). We additionally measured the concurrent depressive symptomatology using the Beck Depression Inventory (BDI; Beck et al., 1961). When recruiting for the study, we ensured we enrolled volunteers with a wide range of anxiety levels to obtain sufficient variability in the cohort.

A subset of participants (N = 86) additionally underwent scanning with a resting-state fMRI paradigm. During the scan, the subjects were instructed to keep their eyes open and focused on a black fixation cross on the white background. The composite anxiety scores were recalculated for the subsample prior to the group analysis. The characteristics of the cohort are provided in Table 1.

**Table 1.** Demographic and psychological characteristics of the participants who constituted the cohorts for the Moral Sentiment and Action Tendencies Task (MSAT), resting-state fMRI (rs-fMRI) and the guilt recollection task (task fMRI).

| Variable | Mean ± standard deviation | Possible range |
| --- | --- | --- |
| Sex | MSAT: 87 females, 53 males<br>rs-fMRI: 54 females, 32 males<br>task fMRI: 49 females, 31 males | not applicable |
| Age | MSAT: 22.56 ± 4.07<br>rs-fMRI: 22.45 ± 3.97<br>task fMRI: 23.80 ± 4.62 | not applicable |
| Trait-anxiety | MSAT: 23.31 ± 12.30<br>rs-fMRI: 23.78 ± 12.89<br>task fMRI: 23.36 ± 12.24 | 0-60 |
| Sensitivity to punishment | MSAT: 12.66 ± 5.94<br>rs-fMRI: 12.60 ± 5.73<br>task fMRI: 12.55 ± 5.76 | 0-24 |
| Behavioral inhibition system | MSAT: 22.19 ± 4.04<br>rs-fMRI: 21.72 ± 4.18<br>task fMRI: 22.43 ± 3.79 | 7-28 |
| Depressive symptoms (BDI) | MSAT: 13.35 ± 10.85<br>rs-fMRI: 13.06 ± 11.50<br>task fMRI: 13.08 ± 9.45 | 0-63 |

Abbreviations: BDI, Beck Depression Inventory.

#### 2.1.2. Task description

The original task was developed in English and translated into Spanish by three native Spanish-speaking members of the research team. When comparing the three independent translations, 9 out of the 28 adjectives did not fully align across translators. These discrepancies were discussed collectively, and a consensus version was established. A certified Spanish–English translator subsequently reviewed the items, confirming close semantic equivalence for 19 of the 28 adjectives, while noting meaningful shifts in nuance for the remaining nine (see Supplementary Table 1). Therefore, the Spanish version should be considered an adapted version rather than strictly equivalent translation, and findings should not be directly compared with studies using the original English materials. Importantly, however, all translated adjectives retained clearly negative, morally or socially disapproving connotations, thereby preserving functional equivalence with respect to the task’s objective of inducing negative moral evaluations. The original English version has been made available online by the authors (https://translational-cognitive-neuroscience.org/test-materials).

In the current study, we used the computerised version of the paradigm. It was administered to the participants during a behavioural experimental session taking place in the week preceding the neuroimaging data acquisition. The task began with a screen asking participants to provide their name and the name of their same-sex best friend with whom they had no family or romantic bonds. Subsequently, they answered a question concerning how close they felt to their friend using a Likert-scale with 7 discrete levels (1 - not at all, 7 - very close). Only participants reporting feeling sufficiently close to their friends (i.e. ≥ 4) were included in the experiment. The value was missing for 1 subject, and 7 participants indicated not feeling sufficiently close to their friend, yielding the aforementioned final sample of 140 individuals.

In the main part of the task, participants were asked to imagine 54 hypothetical scenarios (27 behaviours repeated across 2 agency conditions) in which they (self-agency) or their friend (other-agency) were behaving counter to social and moral values, with the other person being the recipient of the action (e.g. “Michal is being bossy towards Marcos”). For each of such scenarios, subjects were asked how strongly they would blame themselves and their friends in the described situation (Likert scale ranging from 1 (not at all) to 7 (a lot)). Additionally, they were asked what emotion they would feel most strongly with the available choices being four self-blaming emotions (i.e. guilt, shame, contempt/disgust towards self and indignation/anger towards self), two other-blaming emotions (i.e. contempt/disgust towards the friend and indignation/anger towards the friend), as well as no/other emotion. At the end of the trial, the subjects were presented with a choice of 7 actions that they would most likely undertake in the particular scenario: feeling like verbally or physically attacking/punishing themselves, feeling like creating distance from themselves, feeling like hiding, feeling like apologising/fixing what they have done, feeling like verbally or physically attacking/punishing their friend, feel like creating distance from their best friend, as well as no/other action.

### 2.2. fMRI guilt recollection task

#### 2.2.1. Participants

The guilt recollection task in the fMRI setting was performed by 80 individuals, with the majority (N = 71) also forming a part of the MSAT cohort. The composite anxiety scores for the sample were calculated anew, using the same approach as described previously (section 2.1.1.). The characteristics of the cohort are provided in Table 1 (section 2.1.1.).

#### 2.2.2. Task description

During recruitment for the study, participants were asked to provide 7 instances of memories they felt guilty about and 7 instances of emotionally neutral past events. The choice of personally-relevant situations rather than hypothetical guilt-evoking scenarios was guided by the findings that guilt recollection induces higher activity in the relevant brain circuitry (Mclatchie et al., 2016). To ensure the participants’ comfort, the exact details of personal events remained unknown to the experimenters. For each memory, the subjects provided 5 cues, up to 3 words each, that would enable them to identify it afterwards. To make the content of the cues comparable within and across the individuals, we required at least one cue pertaining to the involved people, an associated abstract or concrete object and the location of the event. For both guilt and neutral memories, participants rated how guilty they felt about the particular event and to what extent they felt they had violated the socially accepted norms of behaviour (Likert scale ranging from 1 (not at all) to 7 (very much)). In the case of the neutral events, to ensure their lack of emotional valence, an additional question was asked probing feelings associated with the particular memory. The answers were provided with a Likert scale ranging from −3 to 3, i.e. from extremely negative to extremely positive emotions. For the subsequent part of the experiment, 5 memories from each category were chosen. The inclusion criteria were guilt ratings ≥ 4 for the guilt memories, and emotional valence ratings within the −1 to 1 range for the neutral memories.

In a behavioural experimental session during the week preceding the fMRI scanning, the participants filled in a survey pertaining to the chosen stimuli. In some cases the exact wording of the cues was slightly altered by the experimenters for brevity, and so the participants were asked to confirm their ability to correctly identify the memories. They were also prompted to rate how vividly they could relive emotions associated with each event (Likert scale from 1, not vividly at all, to 4, quite vividly). Additionally, three more questions with prespecified answers concerning the type of location (outdoor, indoor, both), number of people involved (0-2, 3-5, > 5), and the time that had passed since each event took place (≤ 1 year, 2-5 years, > 5 years) were asked. The associated information served as a part of the task performed in the scanner, enabling us to test whether the correct memory was retrieved.

Prior to entering the scanner, the participants were familiarised with the task procedure. To avoid any repetition suppression effects, the stimuli presented during the familiarisation run were different from the participant-specific memories. The guilt recollection task was administered with PsychoPy (version 2022.2.5; Peirce et al., 2019), and was presented to the participants using MRI-compatible goggles (VisuaStim Digital, Resonance Technology Inc.). The behavioural responses were recorded with the right-hand device from the MRI-compatible four-key button box set.

The schematic representation of the guilt recollection task is presented in Figure 1. Each trial began with a 10 s period during which cues related to specific memories were presented. The participants were instructed to ‘relive’ the associated emotions upon recognising the event. In the subsequent stage, the subjects would have 4 s to answer a question regarding the location, socialness or age of the memory. At the end of each trial, the participants performed from 3 to 5 repetitions of a simple magnitude judgment task. Participants had to choose which of the two numbers presented at the bottom of the screen was closer to the target number presented above. The number of arithmetic task repetitions was counterbalanced across the run.

**Figure 1.**
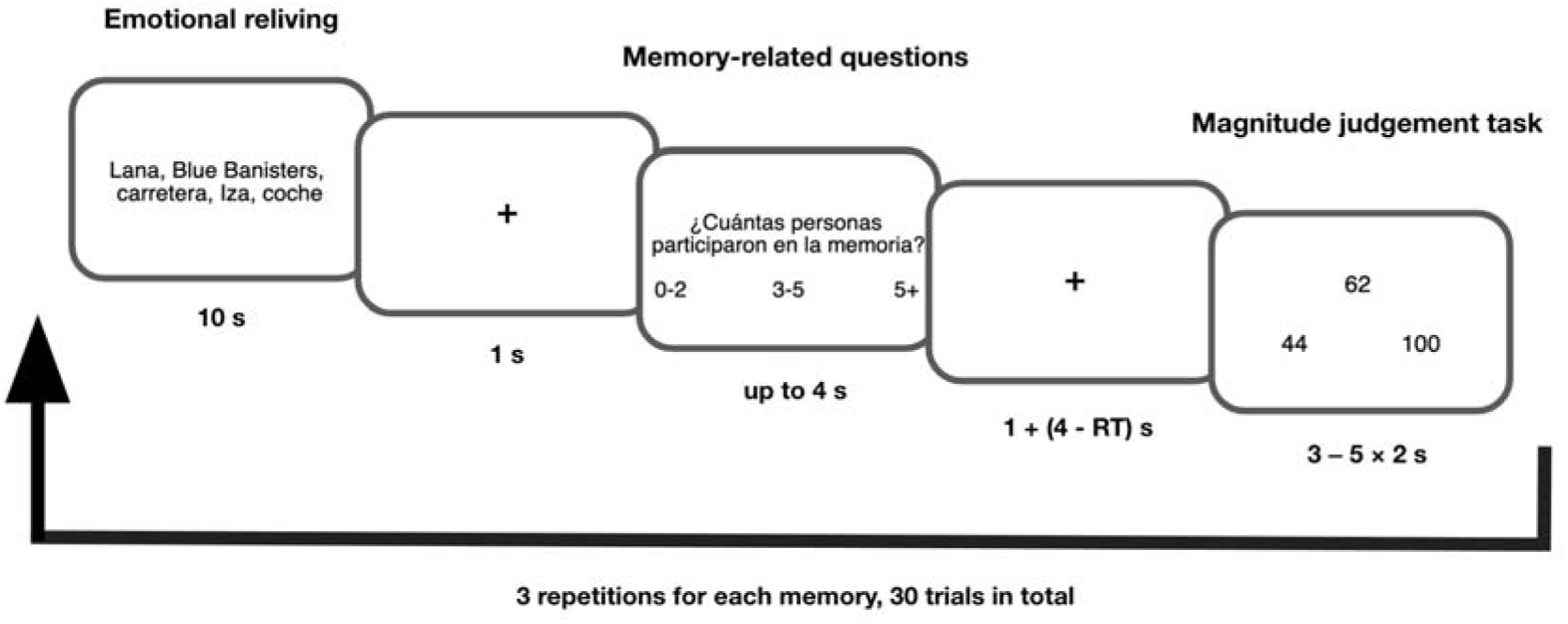
Schematic representation of the guilt recollection task. Abbreviations: RT, reaction time.

We included the memory-related questions and the magnitude judgement task to shift attention of participants from internally- to externally-focused cognition, reducing activity in the socio-emotional brain circuitry before the next trial. The magnitude judgement tasks also served as a non-semantic baseline for the imaging contrast (Binney et al., 2016). For a similar purpose, the guilt and neutral memories were presented in a pseudo-randomised order with not more than two examples of the same condition presented consecutively. All the memories appeared 3 times, each time followed by a question concerning a different aspect of the event, yielding 30 trials in total (13 min).

After the scanning session, participants were asked whether they could relive emotions associated with each memory upon all the repetitions and how vivid the experience had been. In the case of difficulties performing the task, the indicated trials were excluded from the neuroimaging analyses. Additionally, the approach-avoidance motivation related to each event was measured by inquiring whether the participants felt like distancing themselves from the involved people, objects and places, or approaching them (Likert scale, ranging from −3 to 3).

### 2.3. MRI data acquisition and processing

#### 2.3.1. MRI data acquisition

Neuroimaging data was collected with a 3T General Electrics SIGNA Architect scanner. For high-resolution structural images, T1 MP-RAGE sequence was used. The functional T2*-weighted data during the guilt recollection task was collected with a multiband gradient echo-planar imaging sequence (50 axial slices collected in the interleaved order; 2.5 mm isotropic voxel; repetition time = 2 s; echo time = 20.4 ms; flip angle = 90°; multiband acceleration factor = 2; left-right phase-encoding direction). To account for the non-steady state of the magnetic field, 10 volumes were added at the beginning of the sequence, yielding the total scanning time of 13 min 20 s. The decision to use the left-right vs. anterior-posterior phase-encoding direction for the task sequence was guided by higher temporal signal-to-noise ratio (tSNR) observed in the brain areas of interest during the piloting stage, in line with the previous findings suggesting that phase encoding direction matters (Cao et al., 2023; Wang et al., 2023; Halai et al., 2014).

The resting-state fMRI data was collected using the same parameters as described above, the only differences being the use of standard anterior-posterior phase-encoding direction and acquisition of 185 volumes (6 min 10 s). The resting-state data was always acquired between the anatomical and task sequences to avoid the latter’s confounding influence (Li et al., 2016). The task and resting-state fMRI data were both characterised by good signal coverage in the areas of interest to the current study, as evidenced by reliable tSNR levels (see *Data availability* section).

#### 2.3.2. Processing of fMRI data from the guilt recollection task

Preprocessing of the data was conducted using AFNI (Cox, 1996) and FSL (Jenkinson et al., 2012) packages. The decision to use a custom pipeline for the task fMRI data, as compared to the more standardised approach of fMRIprep (Esteban et al., 2019), was guided by suboptimal performance of the latter tool for distortion correction of the images acquired with the current T2*-weighted sequence (i.e. left-right phase-encoding).

Anatomical images were skull-stripped, warped to the MNI space, and segmented to obtain individual cerebrospinal fluid (CSF) masks. The preprocessing of the task data began with deleting the first 5 volumes to allow for signal equilibration. Subsequently, the matrix for motion correction was obtained, and the data was despiked. Then, matrices for susceptibility distortion correction (derived using two separate acquisitions with the opposite phase-encoding directions) and co-registration to the anatomical image were calculated. The spatial transformation of the functional data was done in one step, combining the matrices for motion and distortion correction, coregistration and normalisation to the MNI space.

Following smoothing with a 5 mm Gaussian kernel, the subject-level analysis was performed. The models were run with the pre-whitening option to account for temporal autocorrelations in the data. The nuisance regression included: motion censoring (with a separate regressor for each volume with ≥ 0.5 mm framewise displacement), 12 motion parameters (6 demeaned originals and their first-order derivatives), the CSF time course (physiological noise), and 6 polynomials (low-frequency drifts; determined with the “1 + int(D/150)” equation, where D is the sequence duration in s). The brain activity during each part of the paradigm, i.e. emotional reliving, answering the memory-related questions and magnitude judgement task, was modeled separately by convolving box-car responses with the canonical hemodynamic response function (HRF). The individual brain activity maps for the contrast of interest were generated by subtracting activation patterns associated with the emotional reliving of the guilt and neutral memories for the full epoch length, i.e. 10 s.

Two seeds were chosen for the psychophysiological interactions (PPI) analysis. The left (MNI: −50, 4, −10) and right sATL (MNI: 57, −3, −6) were defined as spheres with a 6 mm radius (Green et al., 2012) centered at the peak coordinates from previous neuroimaging studies on self-blaming (Green et al., 2012; Gifuni et al., 2017; Zareba et al., 2024). The PPI analyses excluded motion censoring to avoid time series discontinuity. To create PPI regressors, mean time courses of seeds were extracted from the residual files of the recalculated task activity-related regression, resulting in the physiological regressors. Following deconvolution, PPI interaction terms were calculated by multiplying the time courses of neural activity with those of psychological task manipulations (i.e. reliving guilt vs. neutral memories) and reconvolving them with HRF. To generate whole-brain maps for the task-dependent connectivity effects, the subject-level analyses were rerun, this time including the physiological and PPI terms.

#### 2.3.3. Processing of resting-state fMRI data

The resting-state data was preprocessed with fMRIprep version 23.2.1. (Esteban et al., 2018), based on Nipype 1.8. (Gorgolewski et al., 2011). In summary, the T1-weighted image was corrected for intensity non-uniformity, skull-stripped, segmented and normalised to the MNI space. The resting-state data was motion, fieldmap and slice-timing corrected, followed by coregistration to the anatomical image and normalisation. To allow for signal equilibration, the first 5 volumes of the functional run were discarded. Such fMRI data was then processed with AFNI (Cox, 1996), where it was smoothed with a 5 mm Gaussian kernel. This step was followed by detrending, denoising and band-pass filtering (0.01-0.1 Hz frequency range), all applied simultaneously. The nuisance regressors included the mean CSF and white matter time courses, the original 6 motion parameters and their first-order derivatives. In the case of the fractional amplitude of low frequency fluctuations (fALFF) calculations, the polynomial detrending was not applied, following recent guidelines (Woletz et al., 2019).

Four types of measurements were derived from the data. Firstly, to probe the magnitude of resting-state activity, ALFF and fALFF were calculated. ALFF was defined as the absolute signal power within the 0.01-0.1 Hz frequency range, while fALFF was calculated as the ratio of ALFF to the total signal power across the entire frequency spectrum. While ALFF has higher test-retest reliability (Zuo et al., 2010), fALFF is more independent from the physiological noise (Zou et al., 2008), rendering the two approaches complementary. Both metrics were mean-normalised within each subject prior to the group analysis.

Secondly, to assess the resting-state functional connectivity within the circuitry of interest, two additional types of measures were calculated. To study the coupling of sATL in detail, we averaged the time courses of the low frequency blood-oxygen-level-dependent (BOLD) signal in the left and right hemisphere ROIs separately, correlated them with the time series of all the other voxels in the brain, and normalised the distribution of the resulting Pearson correlation coefficients using Fisher’s z transform. In turn, to obtain the information on how all the brain regions associated with guilt processing were generally integrated within the relevant circuitry, we first masked each participant’s preprocessed image using the results from the guilt reliving-related one sample t-test, and subsequently calculated global connectivity, i.e. the average correlation coefficient of each voxel’s time series with the rest of the included brain areas.

### 2.4. Primary statistical analyses

The main objectives of the current study were twofold. Firstly, we aimed to investigate how individual anxiety levels were related to the strength of self- and other-blaming emotions, proneness to experiencing particular instances of these, and the corresponding action tendencies, as measured with the MSAT task. Secondly, we sought to characterise how anxiety was associated with brain activity and connectivity patterns during re-experiencing autobiographical guilt memories.

#### 2.4.1. MSAT task

All the described analyses were performed in R (version 4.2.1). The strength of self- and other-blaming emotions experienced by participants were averaged across 27 scenarios in each agency condition. Similarly, we calculated the proportion of trials per agency during which the subjects would most strongly feel a particular emotion and most likely undertake a given action. Such data was used as the dependent variables for permutation-based repeated measures analyses of covariance (ANCOVAs) run using the *aovperm* function from the *permuco* library (Frossand and Renaud, 2021) with 10000 permutations. The models tested the associations of anxiety and its interactions with agency (within-subject condition), controlling for the confounding effects of age and sex. The decision to use a non-parametric approach for the described analyses was guided by the largely non-normal distribution of the emotion- and action-frequency data. Multiple comparison correction was achieved through false discovery rate (FDR < 0.05) applied across each category (i.e. self- vs. other blaming emotions, experienced emotions and undertaken actions). The effect sizes for anxiety-related results (Cohen’s f) were calculated using the *effectsize* library in R (Ben-Shachar et al., 2020).

Additionally, using the original data from the task, where the choices of particular emotions and actions were coded binarily, we ran two types of logistic regression models (*glmer* function from *lme4* library; Bates et al., 2015), similarly to the work of Duan and colleagues (Duan et al., 2021). Firstly, we examined whether apologising, hiding, creating a distance from oneself, and attacking oneself could be predicted by self-blaming emotions (shame, guilt, self-disgust/contempt, and self-indignation) and anxiety, as well as their interactions, controlling for the main effects of age and sex. Secondly, as creating a distance from a friend and attacking them are likely to be motivated by other-blaming emotions (i.e. contempt/disgust towards the friend and indignation/anger towards the friend), we tested such associations using similar models as in the first case, this time putting other-blaming emotions in the place of self-blaming emotions. For the logistic regression analysis, FDR was applied separately for each predictor, taking into account the number of models in which a given variable or interaction term was included.

#### 2.4.2. Guilt recollection task neuroimaging data

The whole-brain analysis of the task neuroimaging data was performed in AFNI. To identify patterns of brain activity and sATL task-dependent connectivity consistently associated with reliving of guilt vs. neutral memories across the cohort, one sample t-tests were run using *3dttest++*. In turn, associations with anxiety were tested using *3dMVM* models (Chen et al., 2014), with age and sex controlled as covariates. Cluster-level family wise error-rate correction (FWE < 0.05) with voxel-level thresholding at p < 0.001 was used for significance testing.

Furthermore, a region-of-interest (ROI) approach was used to investigate the functioning of the sATL - sgACC circuitry in detail. The left and right sATL ROIs were defined as previously described (section 2.3.2), while the bilateral sgACC ROI was set as a 6 mm radius sphere centered at the x = 0, y = 15, z = −5 MNI coordinates (Moll et al., 2006; Zareba et al., 2024). The mean brain activity of all voxels within each of the three areas, as well as PPI of the left and right sATL with sgACC were used as dependent variables. The linear regression models were run with R’s *lm* function, testing for associations with anxiety and keeping age and sex as covariates. FDR was applied to correct for multiple comparisons.

### 2.5. Supporting statistical analyses

The supporting analyses sought to investigate associations of anxiety with the behavioural measures collected for the guilt recollection task. Furthermore, we aimed to analyse how the strength of individual self-blaming emotions from the MSAT task was related to resting-state brain activity and connectivity.

#### 2.5.1. Guilt recollection task behavioural data

For all participants, we calculated separately for guilt and neutral memories the mean values for the following measurements: the associated feelings of guilt, perceived social code violation, vividness of the recall before and during the scanning session, approach-avoidance motivation, salience, as well as accuracy and mean reaction times (RTs) for the questions pertaining to the memory whereabouts. Salience of individual memories was defined as the absolute value of the reported approach-avoidance. Additionally, we calculated the number of times participants could successfully relive in the scanner each memory type. For all of the aforementioned variables, permutation-based repeated measures ANCOVAs were run using the R’s *aovperm* function from the *permuco* library (Frossand and Renaud, 2021) with 10000 permutations. The models tested the associations with anxiety and its interactions with memory type (within-subject condition), controlling for the confounding effects of age and sex. The same approach was used for testing the differences of vividness across each memory type before and during the task execution. To analyse the associations of anxiety with mean emotional valence of neutral memories, as well as well as accuracy and mean RTs for the magnitude judgement task, permutation-based linear regression models were run with the *lmperm* function of the *permuco* library (Frossand and Renaud, 2021) using 10000 permutations. Akin to the analyses described in the section 2.4.1, the effect sizes for anxiety-related results were estimated with the R’s *effectsize* library (Ben-Shachar et al., 2020). Data of one and three participants was excluded from the analysis of accuracy and mean RTs for, respectively, the memory- and arithmetics-related parts of the task due to the use of incorrect response buttons.

#### 2.5.2. Resting-state fMRI data

The voxel-level analyses for ALFF, fALFF, global connectivity and seed-based coupling of the bilateral sATL ROIs were performed using AFNI’s *3dMVM* (Chen et al., 2014). As explained before, in the MSAT participants rated self-blaming emotions (Likert scale 1 to 7) of hypothetical scenarios in which they (self-agency) or their friend (other-agency) were behaving counter to social and moral values. The neuroimaging models investigated associations with the strength of agency-averaged self-blaming emotions, controlling for age and sex. Cluster-level family wise error-rate correction (FWE < 0.05) with voxel-level thresholding at p < 0.001 was used for significance testing. For significant results, we extracted the mean values of the tested metrics inside each cluster, and subsequently correlated them with the composite anxiety scores.

Furthermore, similarly to the task-based analysis, we used the ROI approach to probe the functioning of sATL in detail, extracting their mean ALFF, fALFF and global connectivity values. Their associations with the strength of self-blaming emotions (MSAT) and anxiety scores were tested using separate linear regression models (R’s *lm* function), controlling for the effects of age and sex. Multiple comparison correction was achieved using FDR. The decision not to include the bilateral sgACC in the analyses was guided by the relatively lower tSNR observed in this area with the resting-state sequence, as compared with the task-related data acquired with the left-right phase encoding direction.

### 2.6. Associations with concurrent depressive symptomatology

Last but not least, negative emotional biases are not specific to increased levels of anxiety, additionally forming the core part of depressive symptomatology. Therefore, for all the measures significantly associated with the composite anxiety scores we repeated the analyses, this time using the depressive symptoms measured with BDI (Beck et al., 1961).

## 3. Results

### 3.1. Primary results

#### 3.1.1. MSAT task

The first set of analyses (described in section 2.4.1) tested in the full sample of 140 subjects whether self-blaming emotions and action tendencies measured in the MSAT task differed depending on individual anxiety levels and their interactions with agency. The results confirmed the stronger self-blaming emotions in highly anxious individuals (F = 22.02, p_FDR_ < 0.001, Cohen’s f = 0.40, 95% confidence intervals (CI) [0.23; 0.58]; Figure 2A), which was paralleled by only nominally significant association with a higher prevalence of the guilt feelings (F = 5.85, p_uncorr._ = 0.016, p_FDR_ = 0.114, Cohen’s f = 0.21, 95% CI [0.03; 0.38]; Supplementary Figure 1A). As for the action tendencies, the repeated measures ANCOVA additionally linked elevated anxiety levels with more self-attacking behaviours (F = 7.23, p_FDR_ = 0.046, Cohen’s f = 0.23, 95% CI [0.06; 0.40]; Figure 2B). Contrary to our expectations, however, we did not see significant interactions between anxiety and agency condition for any of the tested measures.

**Figure 2.**
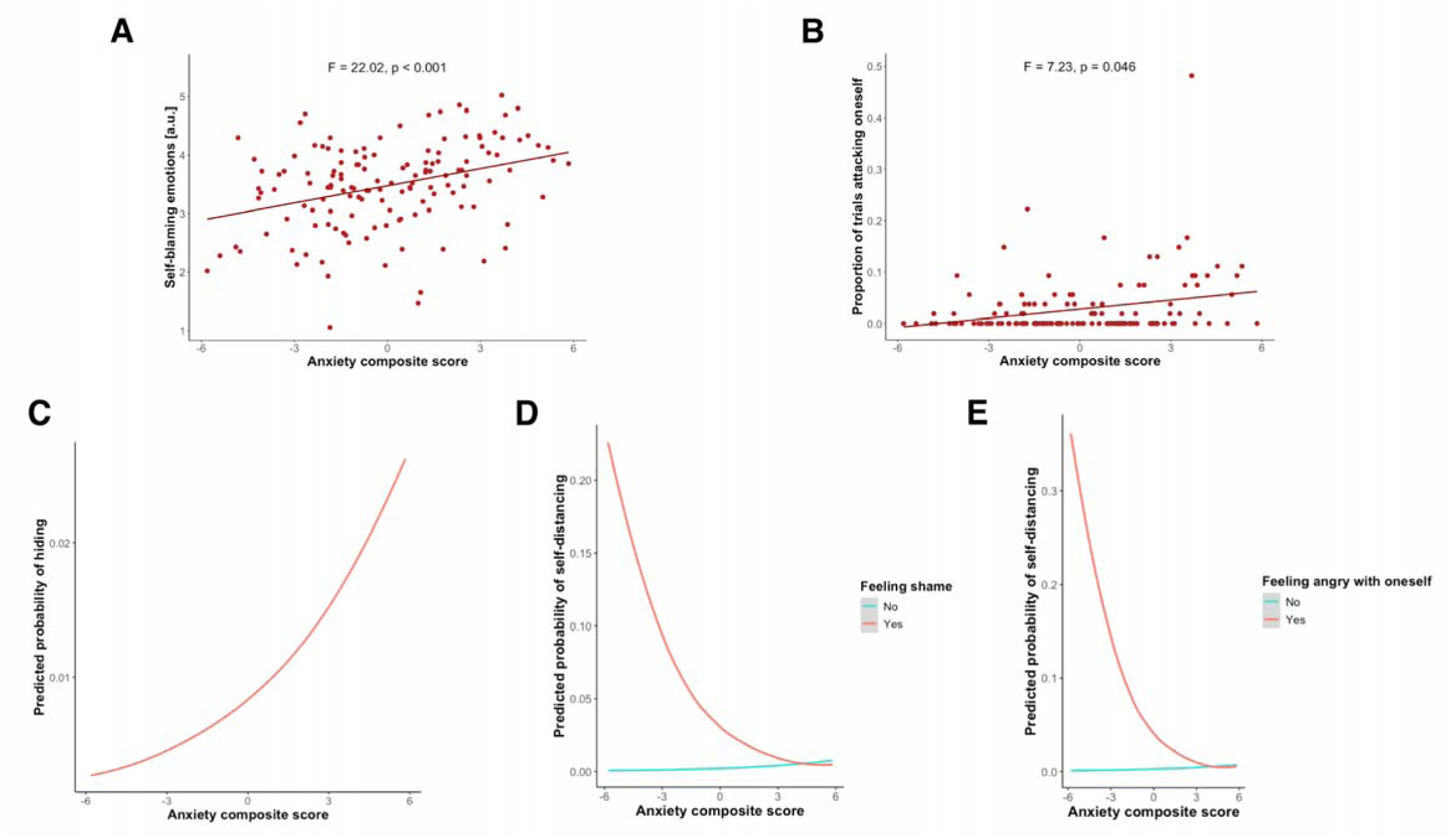
Associations of anxiety levels with self-blaming emotions and action tendencies from the Moral Sentiment and Action Tendencies Task (MSAT). Higher anxiety was related to elevated (A) intensity of self-blaming emotions, (B) self-attacking behaviour and (C) probability of hiding, regardless of experienced emotions. When feeling negative emotions towards oneself, such as (D) shame or (E) anger, highly anxious individuals were also less likely to create a distance from themselves.

The logistic regression analysis complemented the MSAT results by demonstrating, irrespective of the experienced emotions, higher likelihood of hiding in the more anxious participants (β = 0.24, standard error (SE) = 0.08, p_FDR_ = 0.029; Figure 2C). This corroborates the trend-level association obtained using the former approach (F = 4.33; p_uncorr._ = 0.036; p_FDR_ = 0.085; Supplementary Figure 1C). Importantly, logistic regression enabled us to link individual levels of anxiety with specific patterns of behaviour. Namely, highly anxious participants were less likely to create distance from themselves when experiencing shame (β = −0.61, SE = 0.18, p_FDR_ = 0.002; Figure 2D) or self-anger (β = −0.63, SE = 0.19, p_FDR_ = 0.003; Figure 2E). As such, the current results confirm more intense self-blaming emotions in subclinically anxious individuals (Cândea and Szentagotai-Tătar, 2018), linking it with similar maladaptive action tendencies as those observed in MDD patients (Duan et al., 2021; Duan et al., 2023).

The remaining trend-level associations with anxiety can be found in the Supplementary Figure 1, while the findings for agency and sex are demonstrated in Supplementary Figures 2-4.

#### 3.1.2. Brain activity and connectivity during guilt recollection task

Emotional reliving of guilt, as compared to neutral, memories was associated with increased activity in a widespread network consisting of medial and ventrolateral frontal cortices, anterior and posterior temporal lobes, precuneus, insula, striatum, amygdala, hippocampus, thalamus and cerebellum (Supplementary Figure 5). Despite the involvement of bilateral regions, the activation pattern was visibly left-lateralised, which corroborates the previous meta-analytic evidence (Gifuni et al., 2017). In the entire sample, self-blame processing was also related to higher task-related functional connectivity between the right sATL and the left superior frontal gyrus (T = 4.79, k = 31, MNI: −8, 30, 50; Supplementary Figure 6). No such findings were observed for the coupling of the left sATL.

To provide potential insights as to which neuromodulatory systems might contribute to self-blaming symptomatology, we additionally performed a hypothesis-generating analysis investigating whether the mean guilt processing-evoked activity in our sample was correlated with normative neurochemical densities derived from publicly available atlases (Hansen et al., 2022; Gryglewski et al., 2018). The analyses revealed robust associations for all the tested neuromodulatory systems, i.e. serotonin (5-HT), dopamine (DA), norepinephrine (NE) and oxytocin (OXT), indicating that the discussed neurotransmitters might play a crucial role in shaping the neural dynamics when one experiences strong and negative self-referential emotions (Figure 3). Nevertheless, the reported associations do not account for spatial autocorrelation of the neurochemical maps, and should be therefore treated as purely descriptive. The complete methodological details can be found in the section *Exploratory neurochemical correlational analysis* at the end of the Supplementary Material.

**Figure 3.**
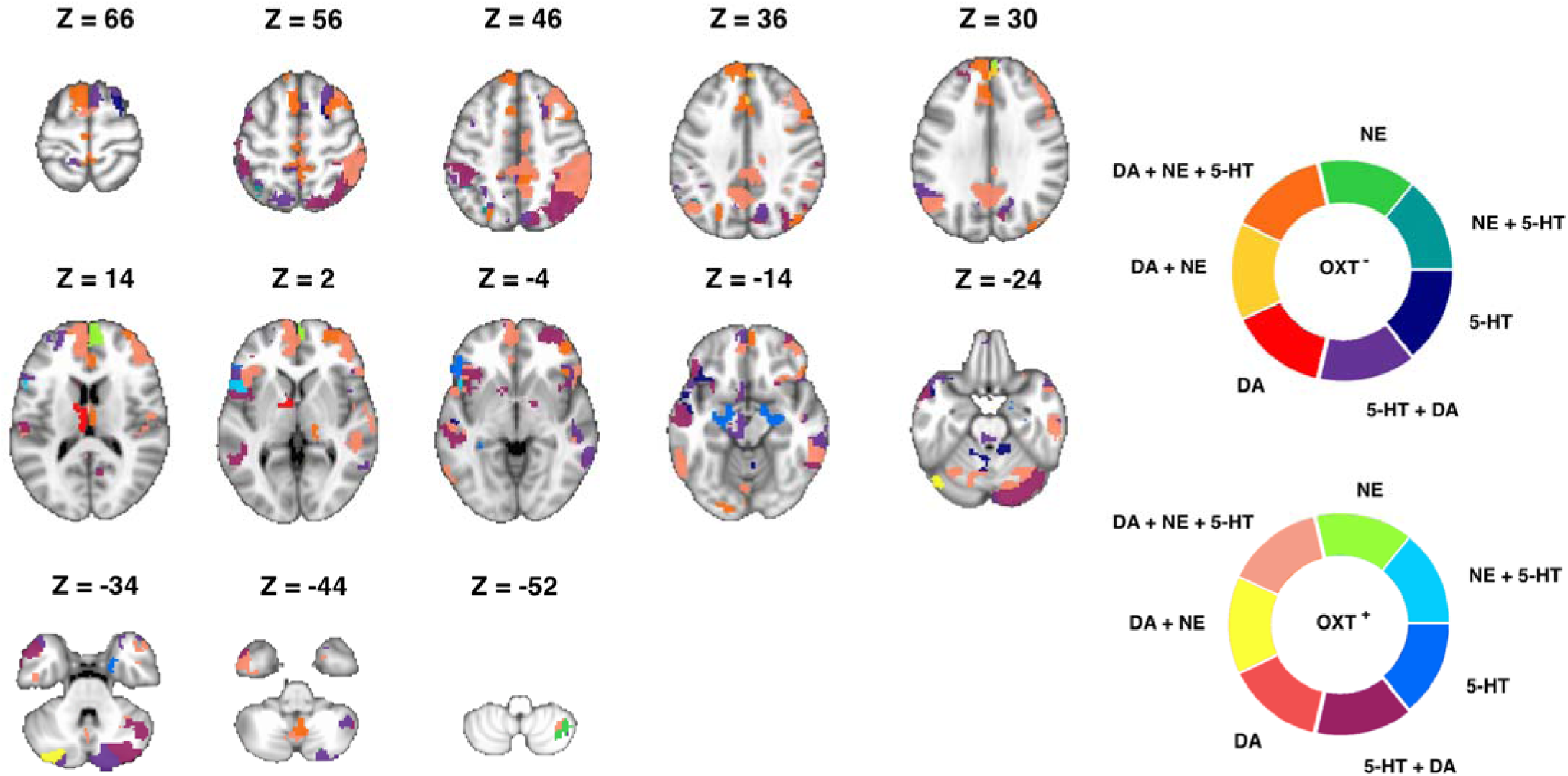
Summary of the regional associations between guilt-related brain activity in our sample and the amounts of neuromodulatory receptors and transporters derived from external normative maps. OXT+ and OXT- stand for, respectively, the presence or lack of associations with the oxytocinergic system. Abbreviations: DA, dopamine; NE, norepinephrine; 5-HT, serotonin.

Regarding the link of the composite anxiety scores with guilt-related activity and connectivity, the whole-brain models revealed no significant findings. In turn, the ROI-level analysis confirmed our hypothesis of increased task-dependent coupling between the left sATL and bilateral sgACC in highly anxious individuals (T = 2.54, p_FDR_ = 0.026, β = 0.07, SE = 0.03). No such associations were found for the right sATL (T = −0.52, p_uncorr._ = 0.604, β = −0.02, SE = 0.03). No effects were similarly observed for the activity of the left sATL (T = −1.39, p_uncorr._ = 0.168, β = −0.05, SE = 0.03), right sATL (T = −0.84, p_uncorr._ = 0.402, β = −0.03, SE = 0.04) or bilateral sgACC (T = −1.86, p_uncorr._ = 0.067, β = −0.06, SE = 0.03).

As explained in section 3.1.1., the MSAT task revealed that increased anxiety was related to higher intensity of self-blaming emotions and maladaptive ways of coping with them. This led us to examine whether the task-dependent functional connectivity between the left sATL and bilateral sgACC was also associated with the discussed behavioural measures using the overlapping sample across the two experiments (N = 71). We followed the same methodology as described previously (section 2.4.1.), with the exception of testing only the influence of guilt in the logistic regression models. A trend-level interaction emerged in which higher task-dependent functional connectivity between the sATL and sgACC was related to lower probability of hiding when one was feeling guilt (β = −0.82, SE = 0.37, p_FDR_ = 0.051; Supplementary Figure 7). Therefore, despite anxious individuals demonstrating a higher probability of socially avoidant behaviours, a guilt-dependent connectivity pattern associated with elevated anxiety was inversely linked to hiding. Such a finding stays in contrast with the expected similarity of the effects, indicating complex associations of behaviour with emotion-specific neural patterns within the studied circuitry. Of note, the discussed task connectivity pattern was unrelated to the strength of self-blaming emotions (r = 0.11, p_uncorr._ = 0.353), the frequency of self-attacking behaviours (ρ = −0.06, p_uncorr._ = 0.628) or the probability of self-distancing when feeling guilt (β = 0.14, SE = 0.56, p_uncorr._ = 0.802).

### 3.2. Supporting results

#### 3.2.1. Behavioural data from the guilt recollection task

The second set of analyses (section 2.5.1.) was focused on the behavioural data associated with the fMRI guilt recollection task. As expected, guilt memories were characterised by higher subjective guilt ratings (F = 5407.89, p < 0.001), perceived social code violation (F = 280.86, p < 0.001) and salience (F = 74.88, p < 0.001), as well as less approach/more avoidant motivation (F = 13.28, p < 0.001) compared to the neutral events (Supplementary Table 2; Supplementary Figure 8). While there were no differences in the vividness of the two types of memories when recalled before (F = 0.84, p = 0.358) or during the scanning session (F = 0.32, p = 0.578), both were characterised by lower ratings during scanning (F = 54.97 and F = 19.15 for, respectively, guilt and neutral memories, both p < 0.001). The vividness ratings nonetheless remained at satisfactorily high level (3.09 ± 0.46 for guilt memories, 3.14 ± 0.75 for neutral memories, with 4 as the maximum score). This confirms that participants could properly relive the associated emotions in both contexts.

The participants were also less accurate (F = 49.45, p < 0.001) and slower (F = 56.47; p < 0.001) when responding to the questions probing whereabouts of the guilt memories. Such an observation suggests increased cognitive engagement as compared to the neutral past events, which is further corroborated by the reports of more difficulty reliving the associated guilt emotions (F = 5.35; p = 0.021). Interestingly, individual anxiety levels positively predicted one’s ability to relive the emotions, regardless of the memory type (F = 4.90, p = 0.031, Cohen’s f = 0.25, 95% CI [0.00; 0.48]; Supplementary Figure 9A). In contrast with our hypotheses, however, anxiety was unrelated to the approach-avoidance motivation (F = 1.96, p = 0.167, Cohen’s f = 0.16, 95% CI [0.00; 0.39]) and salience (F = 2.70, p = 0.106, Cohen’s f = 0.19, 95% CI [0.00; 0.41]) of the guilt memories. Therefore, given the widespread and normal distribution of these two measures (Supplementary Figure 8B-C), they were treated as between-subject factors of interest for an additional exploratory analysis of the task neuroimaging data (section 3.3.1.) that followed the same methodology as described for the anxiety-related results (section 2.4.2.).

The full results of the previously described analyses can be found in Supplementary Table 2. Supplementary Figure 8 demonstrates the observed differences between the guilt and neutral memories, while Supplementary Figure 9B-F depicts associations of their characteristics with sex and age.

#### 3.2.2. Resting-state fMRI analyses

The analysis of the resting-state data indicated that higher resting-state activity (ALFF) in the right temporal pole was related to less intense self-blaming emotions in the MSAT task (F = 26.10, k = 15, MNI: 30, 14, −36). Surprisingly, however, this neural pattern was not related to the composite anxiety scores (r = −0.02, p_uncorr._ = 0.882). The remaining whole-brain models revealed no significant findings for fALFF, global connectivity within the circuitry or the seed-based coupling of the left and right sATL. The ROI-level analysis similarly demonstrated that neither strength of self-blaming emotions, nor the composite anxiety scores were associated with the sATL’s ALFF, fALFF or global connectivity (Supplementary Table 3).

### 3.3. Exploratory fMRI analyses

The whole-brain models for self-blame-related activity revealed no significant associations with the approach-avoidance towards the guilt memories or their salience. In turn, when using the sATL and sgACC ROIs (see section 2.3.2.), we observed higher activity of the left sATL in individuals with greater approach motivation towards their guilt memories (T = 2.69, p_FDR_ = 0.027, β = 0.15, SE = 0.06). No such links were found for either the bilateral sgACC (T = −1.28, p_uncorr._ = 0.203, β = −0.08, SE = 0.06) or the right sATL (T = −0.73, p_uncorr._ = 0.469, β = −0.05, SE = 0.07). Similarly, salience of the guilt memories was unrelated to the activity of the bilateral sgACC (T = −1.16, p_uncorr._ = 0.251, β = −0.18, SE = 0.15), left sATL (T = 1.28, p_uncorr._ = 0.204, β = 0.19, SE = 0.15) or right sATL (T = 1.37, p_uncorr._ = 0.175, β = 0.24, SE = 0.18).

As for the task connectivity, the whole-brain analyses revealed significant findings only for the approach-avoidance domain. Increased aversion towards one’s guilt memories was positively associated with the right sATL’s coupling with ipsilateral paracentral (F = 23.89, k = 43, MNI: 12, −38, 52) and inferior parietal lobules (F = 18.66, k = 27, MNI: 32, −56, 32). The connectivity between the sATL and bilateral sgACC was unrelated to either approach-avoidance towards guilt memories (left sATL: T = 0.46, p_uncorr._ = 0.649, β = 0.02, SE = 0.05; right sATL: T = 1.49, p_uncorr._ = 0.141, β = 0.08, SE = 0.05) or their salience (left sATL: T = 0.07, p_uncorr._ = 0.947, β = 0.01, SE = 0.13; right sATL: T = −0.89, p_uncorr._ = 0.379, β = −0.12, SE = 0.13).

### 3.4 Summary of the neuroimaging results

All the significant neuroimaging results across the previously described analyses are summarised in Figure 4 and Table 2. No associations with age or sex were found.

**Figure 4.**
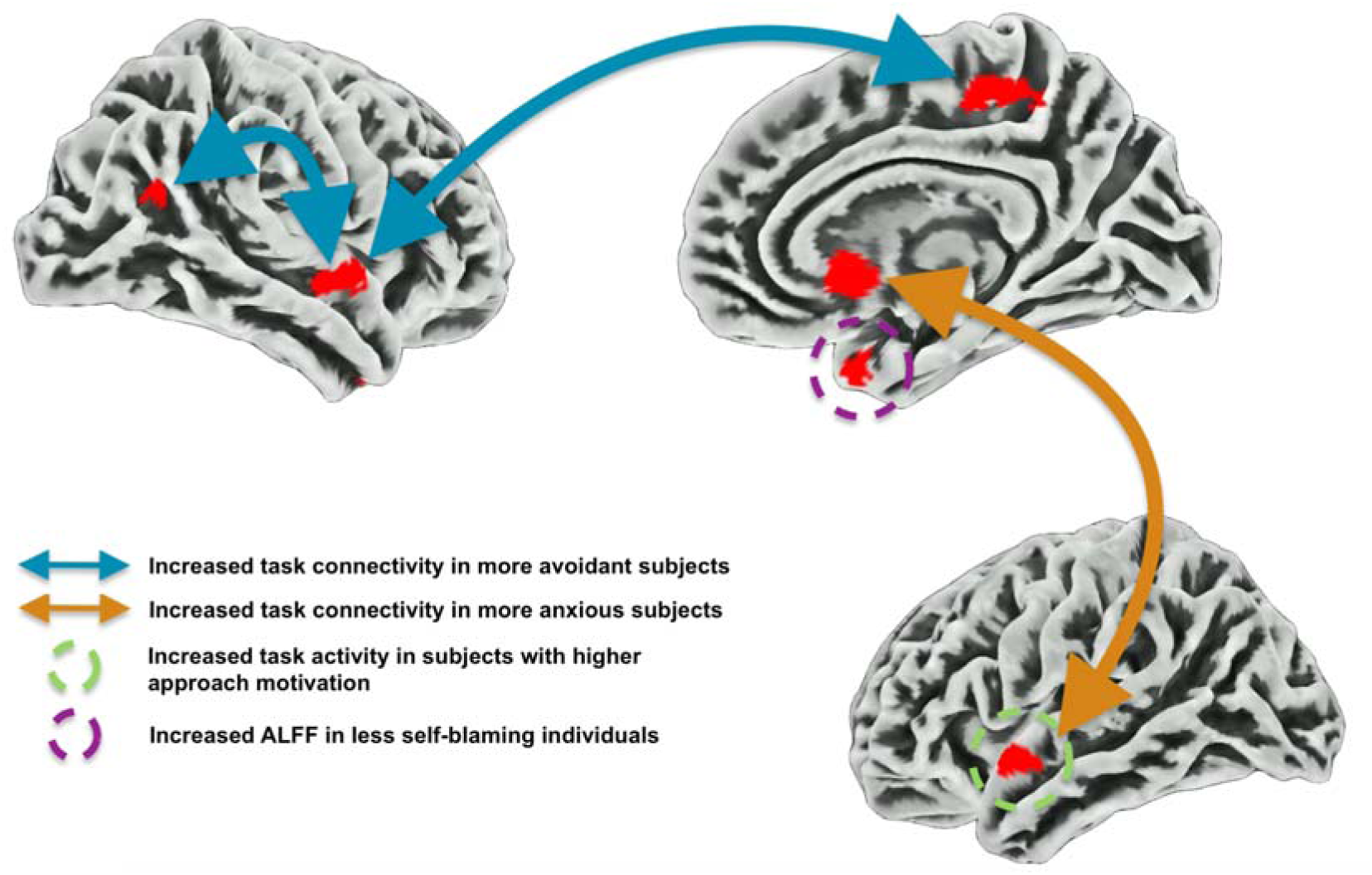
Patterns of brain activity and connectivity associated with subclinical anxiety, approach-avoidance motivation towards guilt memories (guilt recollection task), and the strength of self-blaming emotions from the Moral Sentiment and Action Tendencies task (resting-state). The approach-avoidance dimension was operationalised as a continuum of negative (avoidance) to positive (approach) values. Whole-volume analyses were run for brain activity and the connectivity of the bilateral superior anterior temporal lobes (sATL). Additional region-of-interest (ROI) analyses focused specifically on the sATL’s activity (task and resting-state), and its coupling with bilateral subgenual anterior cingulate cortex (task only). The results pertaining to the higher avoidance motivation and strength of self-blaming emotions survived the cluster-level family-wise error thresholding (p_FWE_ < 0.05), whereas the links with anxiety and higher approach motivation were thresholded at the ROI level using the false discovery rate correction (p_FDR_ < 0.05).

**Table 2.** Summary of the brain activity and connectivity patterns associated with subclinical anxiety, approach-avoidance motivation towards one’s guilt memories, and the strength of self-blaming emotions from the Moral Sentiment and Action Tendencies task. The motivational dimension was operationalised as a continuum of negative (avoidance) to positive (approach) values. Positive relationships therefore indicate increases in neural metrics from the most avoidant to the most approach-motivated individuals. For self-blame-dependent functional connectivity results, the first area represents the seed region.

| Neural measure | Brain region | MNI coordinates | Voxels | Statistics |
| --- | --- | --- | --- | --- |
| **Anxiety** |  |  |  |  |
| Self-blame-dependent functional connectivity | L superior anterior temporal lobe | -50, 4, -10 | 61 | T = 2.54<sup>R</sup> |
|  | B subgenual anterior cingulate cortex | 0, 15, -5 | 61 |  |
| **Approach-avoidance motivation** |  |  |  |  |
| Self-blame processing-related activity | L superior anterior temporal lobe | -50, 4, -10 | 61 | T = 2.69<sup>R</sup> |
| Self-blame-dependent functional connectivity | R superior anterior temporal lobe | 57, -3, -6 | 55 | F = 23.89<sup>W, neg.</sup> |
|  | R paracentral lobule | 12, -38, 52 | 43 |  |
| Self-blame-dependent functional connectivity | R superior anterior temporal lobe | 57, -3, -6 | 55 | F = 18.66<sup>W, neg.</sup> |
|  | R inferior parietal lobule | 32, -56, 32 | 27 |  |
| **Self-blaming emotions intensity** |  |  |  |  |
| Resting-state ALFF | R temporal pole | 30, 14, -36 | 15 | F = 26.10<sup>W, neg.</sup> |

Abbreviations: L, left; R, right; B, bilateral; ALFF, amplitude of low frequency fluctuations.
<sup>R</sup> Region-of-interest analysis, $p_{FDR} < 0.05$ .
<sup>W</sup> Whole-brain analysis, cluster-level $p_{FWE} < 0.05$ .
<sup>neg.</sup> Negative link with the reported psychological measure or an association with more avoidant motivation.

### 3.5. Associations with co-occurring depressive symptoms

Across all the behavioural and neural measures related to anxiety in the previously described analyses, statistically significant associations with depressive symptomatology were only observed for the strength of the self-blaming emotion in the MSAT task (F = 10.40, p_FDR_ = 0.002, Cohen’s f = 0.28, 95% CI [0.10; 0.45]). An additional trend-level link was also present for the decreased probability of self-distancing when feeling angry with oneself (β = −0.09, SE = 0.04, p_FDR_ = 0.066). The summary of the results and their comparison with anxiety-related analyses are presented in the Supplementary Table 4.

## 4. Discussion

The results of the current study demonstrate greater intensity of self-blaming emotions in subclinically anxious individuals, linking it with more pronounced self-attacking and socially avoidant behaviours, irrespective of social agency. Importantly for understanding the aetiology of maladaptive emotion regulation, when experiencing negative emotions about themselves, such as shame and self-anger, highly anxious individuals were less likely to disengage from self-focused thoughts. On the neural level, these findings were paralleled by increased self-blame-dependent functional connectivity between the left sATL and bilateral sgACC in the more anxious participants. Furthermore, stronger self-blaming emotions in the MSAT task were linked with reduced amplitude of resting-state activity in the right temporal pole. Interestingly, despite higher proneness to social avoidance observed across a variety of hypothetical social scenarios in the MSAT task, individual anxiety levels were unrelated to approach-avoidance motivation pertaining to the guilt memories. This aspect of the guilt experiences was nevertheless similarly linked to the sATL circuitry, with higher task activity in the left sATL linked to more approach motivation, and higher self-blame-dependent right sATL connectivity with ipsilateral paracentral and inferior parietal lobules linked to more avoidant feelings.

The presented findings of stronger self-blaming emotions paralleled by maladaptive action tendencies are largely in line with the study’s hypotheses. The identified behavioural patterns are well positioned to explain the symptomatology observed in highly anxious people. The avoidance bias may lead to social withdrawal and isolation, decreasing the likelihood of rewarding social interactions (Silk et al., 2012). Together with self-attacking behaviours, it could reinforce the lower self-esteem typically observed in that group (Fernandes et al., 2022). Moreover, the inability to disengage from self-centered thoughts when feeling negative emotions about oneself might give rise to rumination. Such increased understanding of the self-blaming tendencies in subclinically anxious individuals might therefore prove informative for clinical psychologists and psychotherapeutic efforts.

Interestingly, in the fMRI task, we observed that highly anxious participants reported greater success at reliving the emotional aspects of memories independently of the memory type (guilt vs. neutral). Such a finding does not fully align with the typical accounts of their enhanced encoding of negatively valenced episodic memories, with either intact or diminished encoding of neutral events (Zlomuzica et al., 2014). Nevertheless, recent work demonstrated that elevated anxiety levels are linked to a particularly salient negative mode of episodic memory retrieval, which might create a downstream bias in encoding and subsequent retrieval of otherwise neutral information (Lee and Fernandes, 2017). While the effect was observed across the timescale of minutes, the neutral memories in the current work typically spanned across a week preceding the scanning session. As such, our results suggest that similar processes might also be relevant for episodic memory retrieval on longer, more intermediate timescales. Last but not least, although we hypothesised that higher anxiety levels would be related to increased salience or avoidance towards the guilt memories, we did not observe such associations. This comes as a surprise, given the fact that we observed an avoidance bias in the MSAT task, but could be potentially explained by the inclusion of a limited number of memories uniformly associated with strong feelings of guilt. Such autobiographical events may similarly affect anxious and non-anxious individuals, unlike the broader range of social scenarios tested in the MSAT.

The current study also provides important insights into the alterations within the self-blame processing circuitry that might contribute to these behavioural observations. Confirming our hypothesis, we observed increased self-blame-dependent functional connectivity between the left sATL and bilateral sgACC. This corroborates our previous finding linking the increased functional interconnectedness of the left sATL within the guilt processing network with more frequent self-blaming behaviours (Zareba et al., 2024). The left sATL has been linked to the integration of social and emotional information into semantic knowledge and therefore could be a key region for representing concepts like *guilt* (Binney et al., 2016). This integration could be facilitated via the uncinate fasciculus, a white matter tract which connects the ATL with medial prefrontal regions that include the posterior sgACC (Sakata et al., 2019) and process cues pertaining to self-worth and social affiliation (Will et al., 2017; Zahn, 2025). Therefore, we interpret the increased task-dependent functional connectivity between the two areas as a potential marker of stronger contribution of self-blaming experiences and their associated meaning to the neural computations performed in sgACC. Consequently, this may lead to the feelings of worthlessness and social difficulties observed in anxious individuals (Will et al., 2017; Fernandes et al., 2022). Importantly, similar findings have been reported in MDD patients, albeit for the right sATL, where this neural signature is believed to reflect a fully remitting form of the disorder with higher risk of recurrence but also better chance of remission (Lythe et al., 2015; Fennema et al., 2023; Zahn, 2025).

All in all, the self-blame-related behavioural and neural patterns related to subclinical levels of anxiety in the current study largely resemble observations made previously in MDD patients (Duan et al., 2021; Duan et al., 2023; Lythe et al., 2015; Fennema et al., 2023; Zahn, 2025). Despite that, we found only limited associations with the concurrent depressive symptomatology. As such, these findings are interpretable in line with the accounts of anxious temperament (neuroticism) as a general risk factor for affective disorders, which include both MDD and clinical manifestations of anxiety (Pede et al., 2017; Vinograd et al., 2020). Therefore, it would be of interest to the future longitudinal studies to examine whether self-blaming-related behavioural and neural patterns, like those highlighted by our work, could indeed predict the transition from subclinical levels of anxiety to the more severe symptomatology.

In addition to the findings presented above, we observed increased intensity of self-blaming emotions in participants with lower amplitude of resting-state activity in the right temporal pole. This brain region has been shown to connect social processing with affective cognition (Lambon Ralph et al., 2017). Previous studies have linked the overgeneralisation of information related to self-blame and self-criticism, a feature characteristic for individuals with negative emotional biases (Beck, 1963), to functional disconnection of the right sATL (Green et al., 2013). A body of work suggests, however, that the polar temporal region might also contribute to these observations. Increased activity within this area in females subjected to sexual objectification comments was positively linked to self-esteem and negatively associated with shame proneness (Monachesi et al., 2024). Additionally, larger grey matter volume in that structure was related to seeking and enjoying social relationships (Lebreton et al., 2009), a behaviour that is inextricably linked to self-worth (Harris and Orth, 2020). Our result of increased right temporal pole ALFF being related to less intense self-blaming emotions is therefore in line with the above account. An important methodological distinction with the previous works comes in the fact that in the current study this association was observed during task-free resting-state conditions. Nevertheless, during this period individuals typically engage in retrospective memories or prospective thoughts (Delamillieure et al., 2010), which quite likely requires social semantic and affective processing. Surprisingly, the discussed neural pattern was unrelated to individual anxiety levels, which nevertheless parallels our previous report of only partially overlapping neural correlates between these two phenomena (Zareba et al., 2024). Similarly, the results stemming from this work and previous studies appear to suggest differential involvement of the right vs. left sATL in self-blaming symptomatology observed in MDD and subclinical anxiety (Green et al., 2012; Green et al., 2013; Lythe et al., 2015; Zahn et al., 2019; Jaeckle et al., 2023; Fennema et al., 2023; Zareba et al., 2024). As such, we identify a need for studies more finely delineating the elements of the associated circuitry, with a particular importance placed on identifying the exact cognitive processes governed by specific brain regions and their connections.

Importantly, the current work suggests that approach-avoidance motivation might be one such factor. We observed increased left sATL activity in participants with greater approach towards their guilt memories, and elevated task-dependent coupling of the right sATL with ipsilateral paracentral and inferior parietal lobules in those with stronger avoidance. The potential functional differences between the bilateral sATL appear in line with some previous studies linking increased left sATL activity during social threat perception to lower levels of negative affectivity (Kret et al., 2011), and higher stress-related activity in the right ATL to maladaptive regulation strategies and stress-induced autonomic activity (Golde et al., 2023). Although the associations of increased left- and right-lateralised activity with, respectively, approach and avoidance motivation are predominantly related to the functioning of the frontal brain areas (Kelley et al., 2017), the sATL could be one of the driving forces behind these observations as it communicates the social conceptual information to the frontal regions through feedforward projections of the uncinate fasciculus (Sakata et al., 2019). Importantly for the avoidance-related results, the right sATL is also structurally connected to the ipsilateral paracentral and inferior parietal lobules (Fan et al., 2016). While the increased self-blame-dependent connectivity with the former area might represent embodiment of the avoidance concept (Zhou et al., 2024), in the case of the right inferior parietal lobule the higher coupling might signify stronger integration of negative social feedback (Davis et al., 2023), and its contribution to the emergence of guilt-related avoidance (Gao et al., 2021).

Last but not least, the exploratory neurochemical correlational analysis indicated that DA, NE, 5-HT and OXT systems might be differentially associated with the guilt-evoked activity in the relevant brain networks. Previous psychopharmacological studies demonstrated that manipulating neuromodulatory systems indeed influences guilt and moral behaviour (Kanen et al., 2021; Li and Zu, 2024; Zheng et al., 2024). The current work therefore hints at the brain areas that could potentially mediate these effects. It needs to be stressed, however, that the associations reported in the current study do not account for spatial autocorrelation of the neurochemical maps, limiting the inferences that can be drawn. Furthermore, our understanding of how neuromodulators contribute to the observed BOLD responses is still largely limited (Bruinsma et al., 2018). Despite such limitations, previous works utilising a similar methodology were successful in predicting the effects of acute pharmacological manipulations on brain activity and connectivity (Xu et al., 2024). Therefore, these preliminary results warrant further investigation into the utility of such approaches for personalised psychiatry, where use of patient-specific neuroimaging data could guide the choice of pharmacological treatment.

Regarding the clinical implications of the findings, we also stress the subclinical characteristics of the current cohort. While such populations likely share the neural basis for their symptomatology with the diagnosed individuals (Besteher et al. 2017), we cannot exclude the possibility that these associations may become altered in patients with a longer history of disorder (Agorastos et al., 2020). As such, the reported results require replication in subjects meeting clinical diagnosis. The inclusion of the subclinical sample could also contribute to the lack of hypothesised findings for increased salience and avoidance towards guilt memories in highly anxious individuals. Regarding the motivational aspect of guilt memories, we additionally acknowledge the fact that the cross-sectional design deployed in our study to link it with indices of neural activity is likely characterised by lower statistical power compared to the approach based on within-subject comparisons. Nonetheless, the distribution of individual participants’ data precluded the use of the latter method.

Additionally, the present task-based fMRI study contrasts brain activity during recall of autobiographical guilt and neutral memories. Therefore, the described neural patterns cannot be attributed as selective to self-blaming. Nevertheless, identifying regions that are specific to self-blame was not the aim of the present study as prior research has already advanced these research questions (Eslinger et al., 2021; Zahn, 2025). Instead, our objective was to use self-blaming emotions as a particularly salient example of negative self-referential emotions. Therefore, the future studies are encouraged to investigate whether the neural patterns observed in the current work are potentially selective to self-blaming, for instance by contrasting it with other-blaming conditions (Lythe et al., 2015). Furthermore, it would be beneficial to see whether the altered task-related coupling between the sATL and posterior sgACC is also generalisable to other phenomena implicated in anxious symptomatology, such as social comparisons (Acuña et al., 2025). This could potentially help establish integration of social semantic information with self-worth and social affiliation processing as a domain-general cognitive process of interest for clinical interventions in anxiety patients.

Finally, as explained in the Methods section (2.1.2.), we have used an adapted version of the MSAT task for the behavioural paradigm. The translation of the items into Spanish was not one-on-one as some of the meanings of the stimuli were changed. In addition, it needs to be considered that in Spanish the same term (“culpa”) is commonly used to refer both to guilt and self-blame. This means that the lexical distinction between guilt (referring to the emotion about a specific behaviour) and self-blame (the cognitive attribution of the negative behaviour to the self) is absent in Spanish, posing a limitation to the results obtained with that task.

## Conclusions

In summary, the current study links more intense self-blaming emotions in subclinically anxious individuals with specific maladaptive patterns of behaviour, such as more pronounced self-attacking and hiding, as well as inability to disengage from self-focused cognitions when feeling negative emotions about oneself. On the neural level, these observations were paralleled by increased self-blame-dependent functional connectivity between the left sATL and bilateral sgACC, suggestive of stronger contribution of self-blaming experiences to the self-worth and social affiliation computations performed in the latter area. Importantly, distinct neural characteristics pertaining to the bilateral temporal circuitry were modulated by the strength of participants’ self-blaming emotions and the approach-avoidance motivation towards their guilt memories. Combined, these findings highlight the complex, multidimensional nature of the phenomenon, together with the crucial role the social semantic processing appears to play in it.

## Data availability

The guilt recollection task, the neuroimaging analyses scripts and anonymised data from the MSAT task will be made available on a GitHub repository upon acceptance of the manuscript. (1) Mean tSNR maps from the guilt recollection task and resting-state sequence, (2) statistical maps showing the main group effects and associations with anxiety, approach-avoidance motivation and salience for the guilt recollection paradigm, (3) the parcellation scheme used for investigating links between guilt-related activity and the normative amounts of neuromodulatory transporters and receptors, (4) and the thresholded cross-modality association maps are available in the following NeuroVault repository: https://neurovault.org/collections/21877/.

## CRediT author statement

MRZ: Conceptualization, Methodology, Investigation, Formal analysis, Writing - Original Draft, Visualisation. IGG: Methodology, Resources, Investigation, Formal analysis, Writing - Review & Editing. MIM: Resources, Investigation, Writing - Review & Editing. RB: Conceptualization, Writing - Review & Editing. PH: Conceptualization, Writing - Review & Editing. MV: Supervision, Conceptualisation, Methodology, Investigation, Writing - Review & Editing.

## Acknowledgements

We would like to thank Prof. Roland Zahn, Dr. Suqian Duan and Dr. Diede Fennema of King’s College London for kindly providing us with the MSAT task. We would also like to express our gratitude to Raphael Kaplan for his feedback on the guilt recollection task design, as well as Lidon Marin-Marin, Giacomo Tartaro, Marta Rodriguez Aramendia and Tatiana Davydova for their feedback on the translation of the MSAT task.

## Ethical statement

All the procedures followed were in accordance with the ethical standards of the responsible committee on human experimentation (institutional and national) and with the Declaration of Helsinki (1975), and the applicable revisions at the time that this research was underway. Informed consent to be included in the study was obtained from all the participants. The study protocol was approved by the Universitat Jaume I Ethics Committee (CEISH/07/2022).

