## Supplementary Material for "Subclinical anxiety is associated with reduced self-distancing and enhanced self-blame-related connectivity between anterior temporal and subgenual cingulate cortices"

**Supplementary Table 1.** Translation of the Moral Sentiment and Action Tendencies (MSAT) task items from English to Spanish.

| English (original) | Spanish (translation) | Change of meaning |
|---|---|---|
| is apathetic towards | es antipático con | x |
| is boastful towards | es arrogante con | x |
| is bossy with | es mandón con | |
| is careless towards | es descuidado con | x |
| is competitive with | es competitivo con | |
| is cruel towards | es cruel con | |
| is deceitful towards | es engañoso con | |
| is disagreeable towards | es desagradable con | |
| is dishonest with | es deshonesto con | |
| is firm with | es estricto con | |
| is greedy towards | es avaricioso con | |
| is helpless towards | es dependiente de | x |
| is ignorant towards | es insensible con | x |
| is impatient with | es impaciente con | |
| is jealous towards | está celoso de | |
| is possessive towards | es posesivo con | |
| is prejudiced towards | es exigente con | x |
| is quarrelsome with | es agresivo con | x |
| is rebellious towards | es maleducado con | x |
| is romantic towards | es romántico con | |
| is rough towards | es brusco con | |
| is selfish towards | es egoísta con | |
| is stingy towards | es tacaño con | |
| is temperamental with | es temperamental con | |
| is touchy with | es molesto con | x |
| is tough towards | es duro con | |
| is vain towards | es vanidoso con | |

**Supplementary Table 2.** Associations with the behavioural measures collected in the procedure related to the guilt recollection task. Bolded values indicate significant links.

| **Measure** | **Anxiety test value** | **Anxiety p-value** | **Memory type F-stat** | **Memory type p-value** | **Interaction F-stat** | **Interaction p-value** | **Age test value** | **Age p-value** | **Sex test value** | **Sex p-value** |
|---|---|---|---|---|---|---|---|---|---|---|
| Guilt ratings | 2.15 $^{F}$ | 0.142 | **5407.89** | **0.0001** | 0.99 | 0.327 | **7.33 $^{F}$** | **0.01** | 0.21 $^{F}$ | 0.658 |
| Perceived social code violation | 0.06 $^{F}$ | 0.814 | **280.86** | **0.0001** | 0.90 | 0.366 | **7.16 $^{F}$** | **0.0081** | **6.94 $^{F}$** | **0.011** |
| Emotional valence of neutral memories | 0.17 $^{T}$ | 0.869 | n/a | n/a | n/a | n/a | 0.53 $^{T}$ | 0.603 | 0.72 $^{T}$ | 0.479 |
| Vividness (before the scanning session) | 0.34 $^{F}$ | 0.557 | 0.84 | 0.358 | 2.98 | 0.09 | 0.84 $^{F}$ | 0.361 | 1.99 $^{F}$ | 0.173 |
| Vividness (inside the scanner) | 1.62 $^{F}$ | 0.203 | 0.32 | 0.578 | 0.07 | 0.799 | 2.36 $^{F}$ | 0.128 | 0.12 $^{F}$ | 0.728 |
| Successful reliving of emotions inside the scanner | **4.90 $^{F}$** | **0.031** | **5.35** | **0.021** | 0.23 | 0.625 | 0.03 $^{F}$ | 0.870 | 0.09 $^{F}$ | 0.762 |
| Approach-avoidance motivation | 1.96 $^{F}$ | 0.167 | **13.28** | **0.0006** | 0.37 | 0.544 | 3.23 $^{F}$ | 0.074 | 1.94 $^{F}$ | 0.170 |
| Salience | 2.70 $^{F}$ | 0.106 | **74.88** | **0.0001** | 0.35 | 0.559 | 2.22 $^{F}$ | 0.148 | 0.00 $^{F}$ | 0.956 |
| Difference in approach-avoidance motivation between guilt and neutral memories | -0.81 $^{T}$ | 0.415 | n/a | n/a | n/a | n/a | 0.05 $^{T}$ | 0.959 | 1.72 $^{T}$ | 0.089 |
| Difference in salience between guilt and neutral memories | -0.60 $^{T}$ | 0.549 | n/a | n/a | n/a | n/a | 0.17 $^{T}$ | 0.873 | -0.46 $^{T}$ | 0.651 |
| Accuracy for the memory content-related questions | 0.01 $^{F}$ | 0.909 | **49.45** | **0.0001** | 1.99 | 0.168 | 0.68 $^{F}$ | 0.411 | 0.00 $^{F}$ | 0.991 |
| Mean reaction time for the memory content-related questions | 0.76 $^{F}$ | 0.385 | **56.47** | **0.0001** | 0.63 | 0.432 | **4.26 $^{F}$** | **0.0423** | **5.80 $^{F}$** | **0.0218** |
| Accuracy in the mental arithmetics task | -0.03 $^{T}$ | 0.976 | n/a | n/a | n/a | n/a | 0.45 $^{T}$ | 0.644 | 0.45 $^{T}$ | 0.676 |
| Mean reaction time in the mental arithmetics task | 0.58 $^{T}$ | 0.568 | n/a | n/a | n/a | n/a | -0.50 $^{T}$ | 0.623 | 1.04 $^{T}$ | 0.298 |

Abbreviations: $^{F}$, F-stat; $^{T}$, T-stat; n/a, non-associated.

**Supplementary Table 3.** Associations of the strength of the self-blaming emotions and the composite anxiety scores with resting-state characteristics of the bilateral superior anterior temporal lobes (sATL).

| **Region** | **Neural measures** | **Self-blaming emotions T-stat (p value)** | **Composite anxiety score T-stat (p value)** |
|---|---|---|---|
| L sATL | ALFF | 0.72 (0.473) | 0.51 (0.611) |
| L sATL | fALFF | -1.55 (0.126) | 0.31 (0.756) |
| L sATL | global connectivity | 1.74 (0.086) | 1.88 (0.064) |
| R sATL | ALFF | 0.17 (0.865) | 0.94 (0.349) |
| R sATL | fALFF | -0.88 (0.381) | -0.85 (0.398) |
| R sATL | global connectivity | 1.10 (0.274) | 0.45 (0.651) |

Abbreviations: L, left; R, right; ALFF, amplitude of low frequency fluctuations; fALFF, fractional ALFF.

**Supplementary Table 4.** Associations of the anxiety-related results with the concurrent depressive symptomatology. For the measures where effect sizes are provided as Cohen's f, the values below indicate the 95% confidence intervals (CI). If effect magnitude is provided in the form of β, the value below signifies its standard error (SE). Statistically significant effects for depressive symptomatology are presented in bold, while the trend-level associations are additionally put in italics.

| **Measure** | **Associations with anxiety** | | | **Associations with BDI scores** | | |
|---|---|---|---|---|---|---|
| | **Effect size [95% CI/SE]** | **Statistics** | $p_{FDR}$ | **Effect size [95% CI/SE]** | **Statistics** | $p_{FDR}$ |
| ***MSAT, repeated measures ANCOVA*** | | | | | | |
| Self-blaming emotions intensity | Cohen's f = 0.40 [0.23; 0.58] | F = 22.02 | < 0.001 | Cohen's f = 0.28 [0.10; 0.45] | F = 10.40 | **0.002** |
| Proportion of trials attacking oneself | Cohen's f = 0.23 [0.06; 0.40] | F = 7.23 | 0.046 | Cohen's f = 0.22 [0.04; 0.39] | F = 6.36 | 0.110 |
| ***MSAT, logistic regression*** | | | | | | |
| Predicted probability of hiding | β = 0.24 [0.08] | —- | 0.029 | β = 0.04 [0.02] | —- | 0.135 |
| Predicted probability of self-distancing when feeling shame | β = -0.61 [0.18] | —- | 0.002 | β = -0.06 [0.03] | —- | 0.202 |
| Predicted probability of self-distancing when feeling self-anger | β = -0.63 [0.19] | —- | 0.003 | β = -0.09 [0.04] | —- | ***0.066*** |
| ***Guilt recollection task*** | | | | | | |
| Self-blame-dependent L sATL - B sgACC functional connectivity | β = 0.07 [0.03] | T = 2.54 | 0.026 | β = 0.01 [0.01] | T = 1.25 | 0.424 |
| Successful reliving of emotions inside the scanner (number of memories) | Cohen's f = 0.25 [0.00; 0.48] | F = 4.90 | 0.031 | Cohen's f = 0.03 [0.00; 0.25] | F = 0.09 | 0.774 |

Abbreviations: $p_{FDR}$, false-discovery rate corrected p-value; BDI, Beck Depression Inventory; MSAT, Moral Sentiment and Action Tendencies task; L sATL, left superior anterior temporal lobe; B sgACC, bilateral subgenual anterior cingulate cortex.

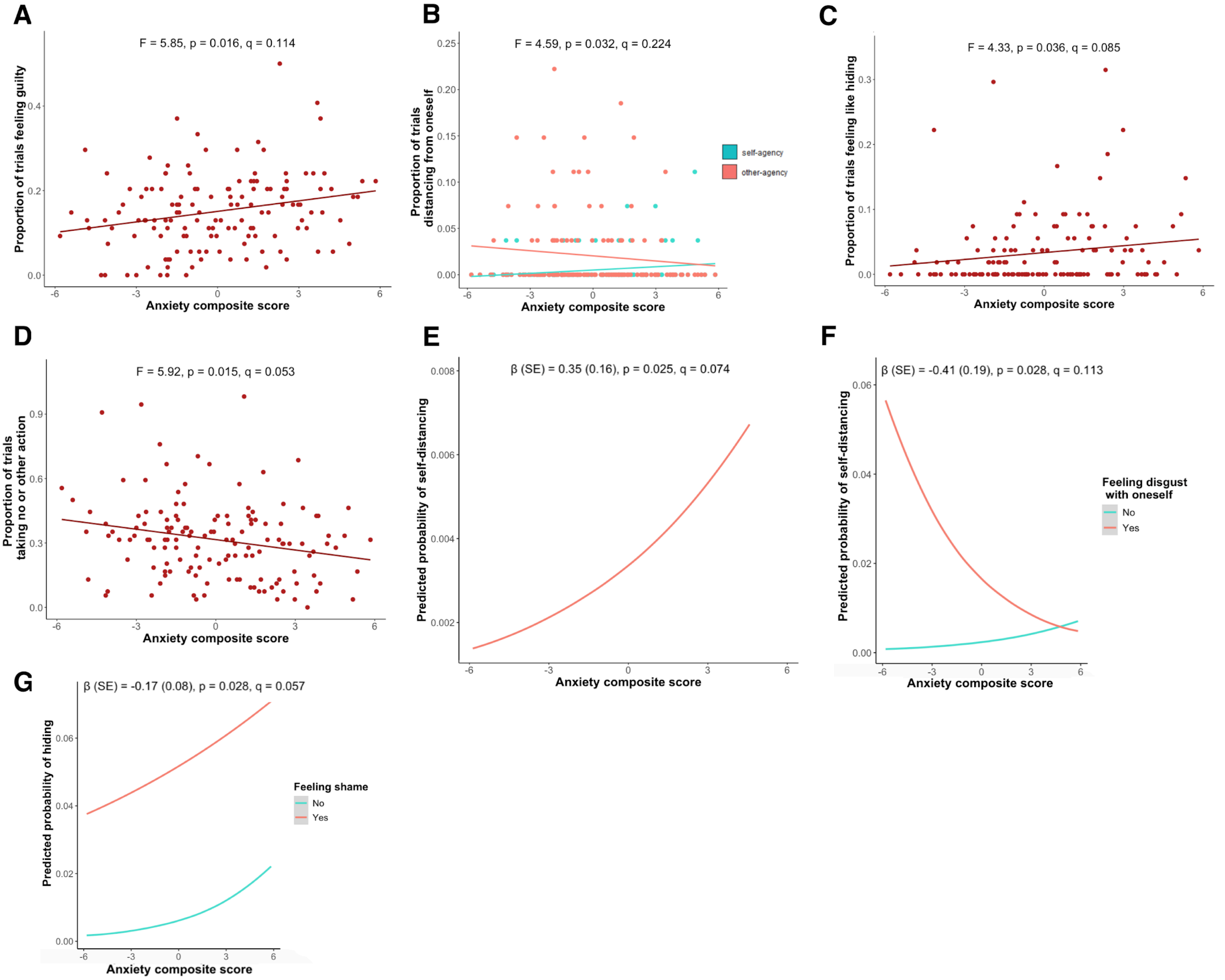


**Supplementary Figure 1.** The trend-level associations of anxiety with the: (A) proportion of trials when participants felt guilty; (B) proportion of trials when participants felt like creating distance from themselves, depending on agency; (C) proportion of trials when participants felt like hiding; (D) proportion of trials when participants would take no or other action; (E) predicted probability of self-distancing, regardless of the felt emotions; (F) predicted probability of self-distancing when feeling disgust towards oneself; (G) predicted probability of hiding when feeling shame. Abbreviations: p, nominal p-value; q, false discovery rate-corrected p-value; SE, standard error.

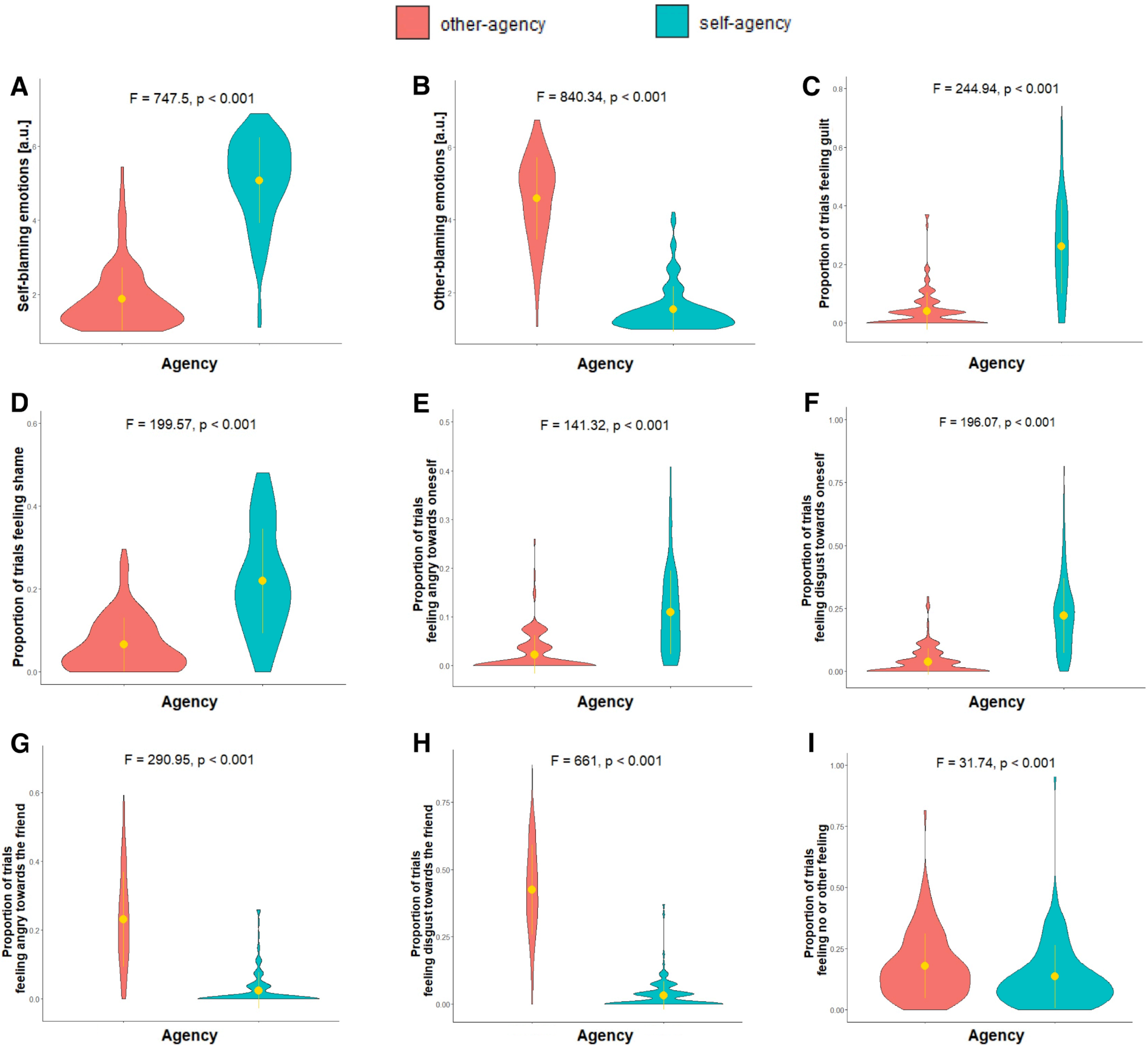


**Supplementary Figure 2.** The effect of agency condition on the self- and other-blaming emotions: (A) strength of self-blaming emotions; (B) strength of other-blaming emotions; (C) proportion of trials feeling guilt; (D) proportion of trials feeling shame; (E) proportion of trials feeling angry towards oneself; (F) proportion of trials feeling disgust towards oneself; (G) proportion of trials feeling angry towards the friend; (H) proportion of trials disgust towards the friend; (I) proportion of trials feeling no or other feeling. The dots and lines represent, respectively, the mean and standard deviation. The statistical significance values were FDR-adjusted.

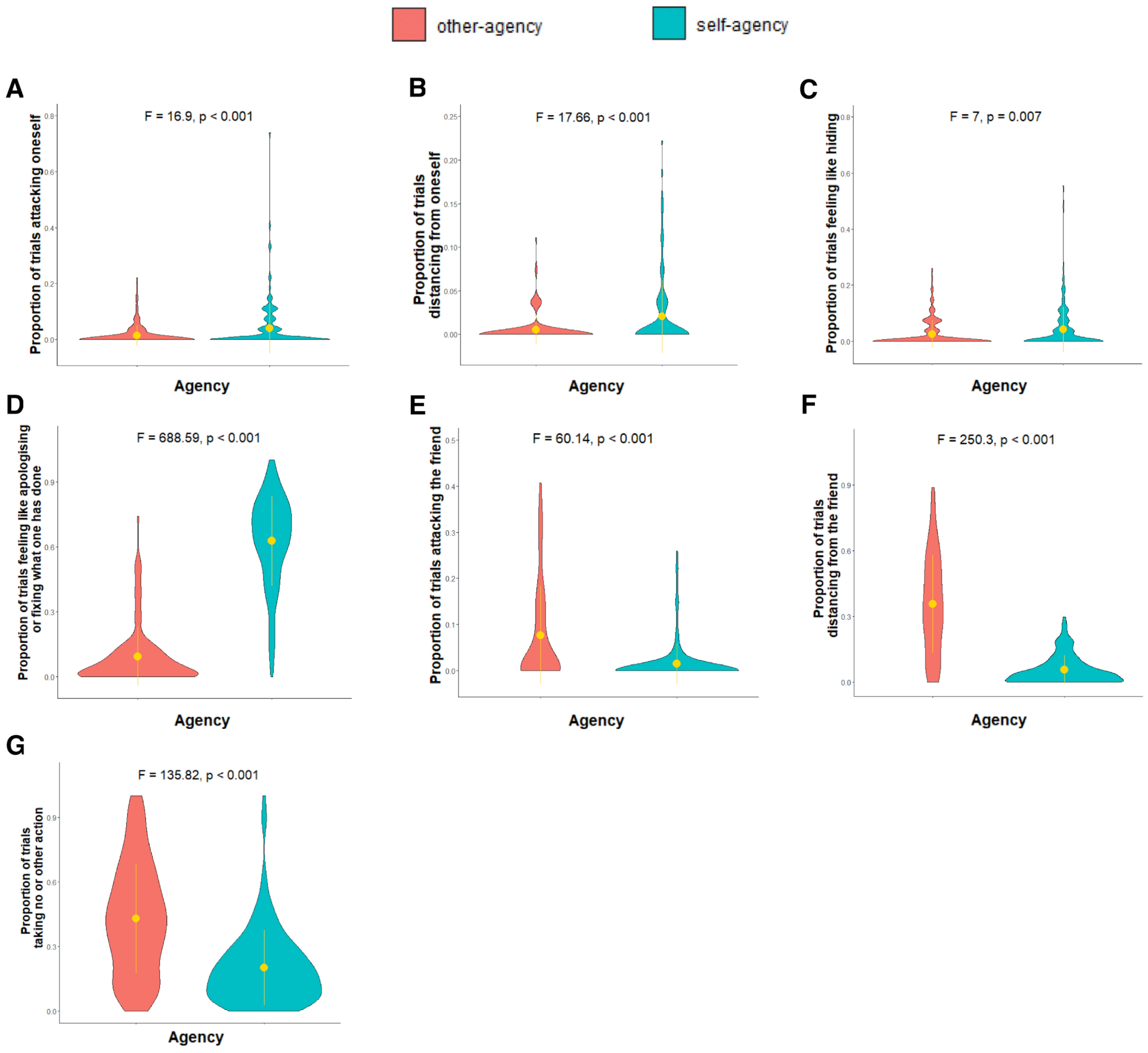


**Supplementary Figure 3.** The effect of agency condition on the action tendencies: (A) feeling like verbally or physically attacking/punishing oneself; (B) feeling like creating distance from oneself; (C) feeling like hiding; (D) feeling like apologising/fixing what one has done; (E) feeling like verbally or physically attacking/punishing the friend; (F) feeling like creating distance from the friend; (G) taking no or other action. The dots and lines represent, respectively, the mean and standard deviation. The statistical significance values were FDR-corrected.

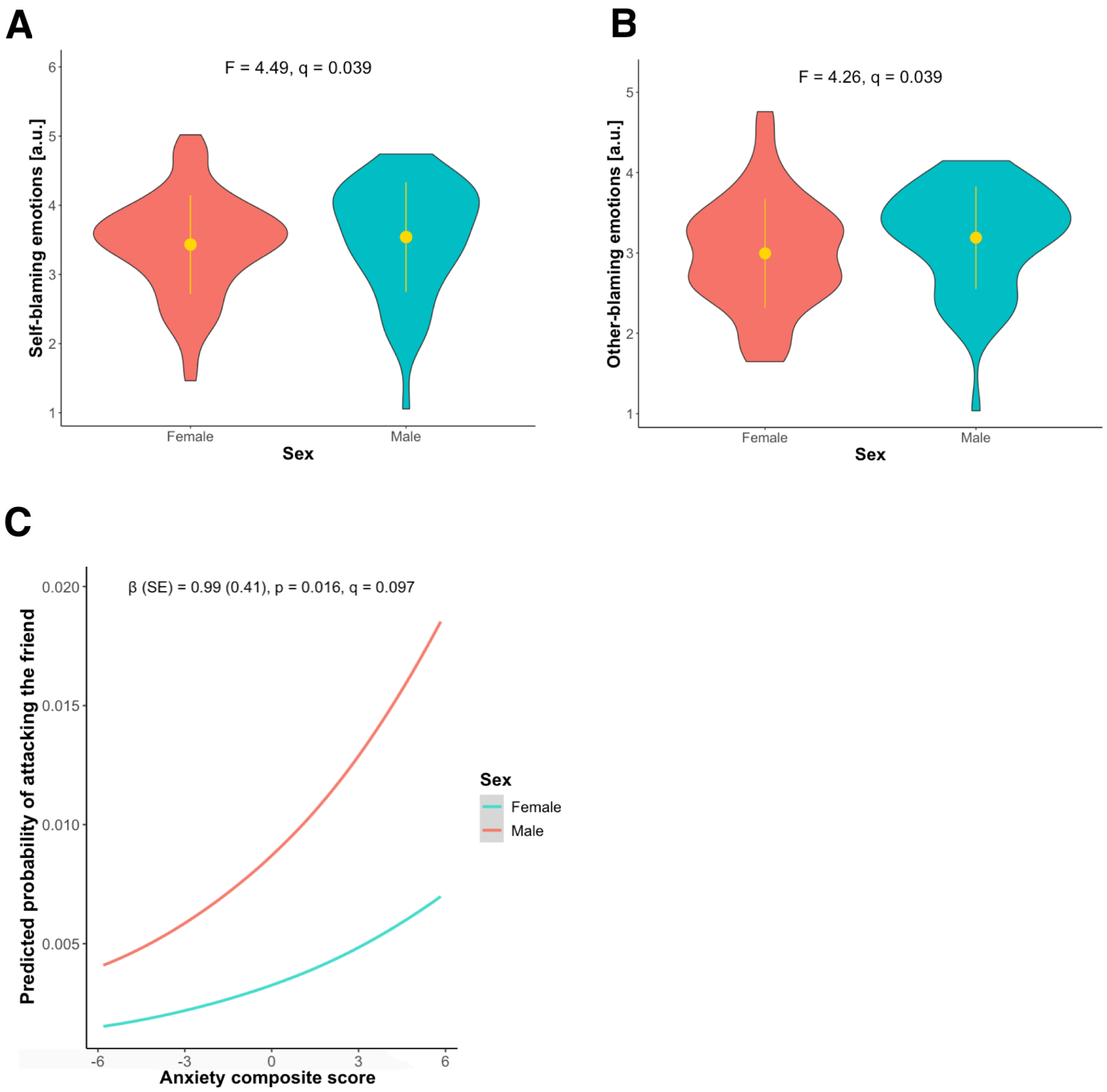


**Supplementary Figure 4.** Male participants reported higher (A) self-blaming emotions, (B) other-blaming emotions, and were also (C) more likely to attack their friends, regardless of the felt emotions. The dots and lines in Figures A and B represent, respectively, the means and standard deviations. Abbreviations: p, nominal p-value; q, false discovery rate-corrected p-value; SE, standard error.

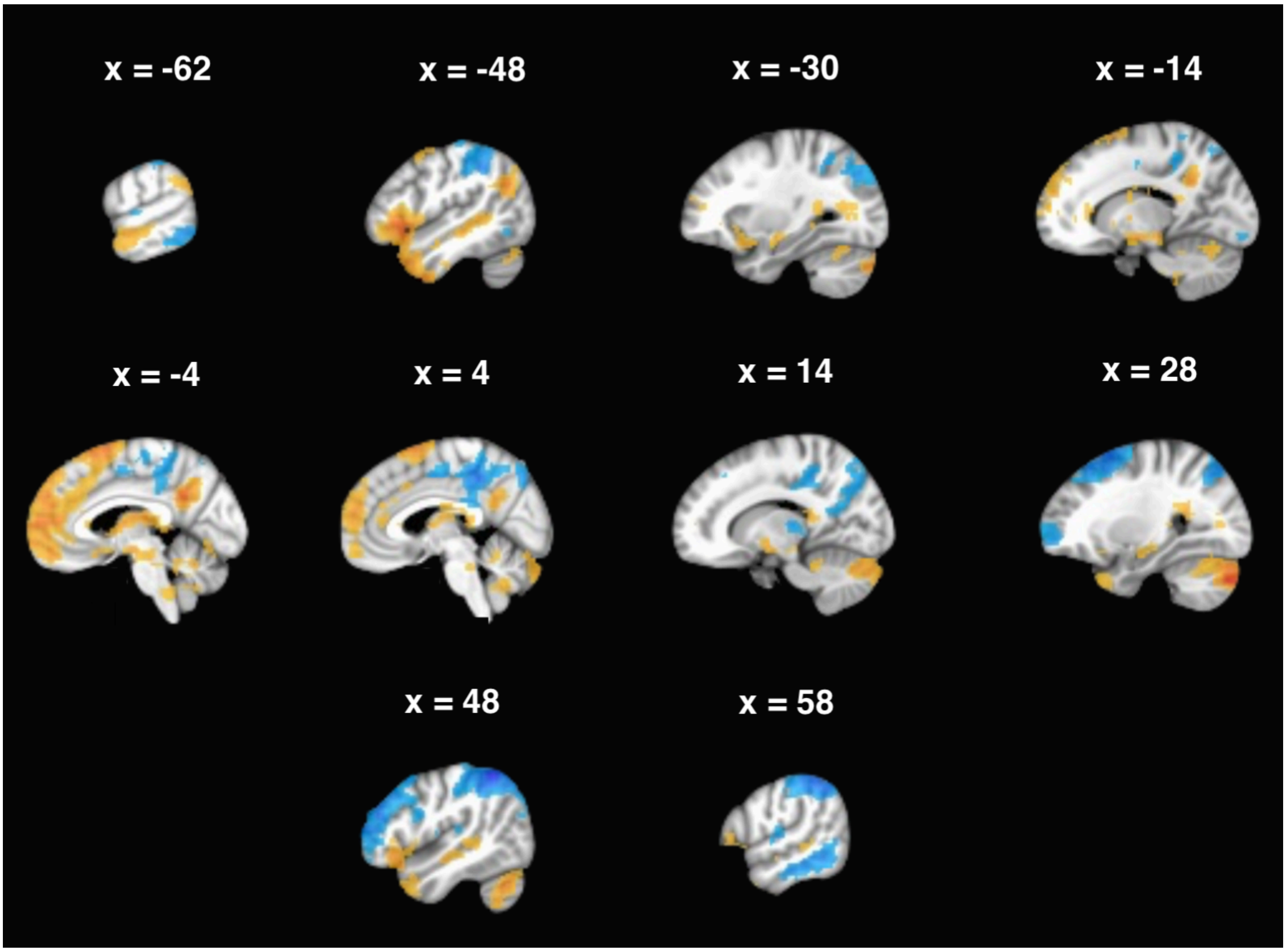


**Supplementary Figure 5.** Brain areas with significantly increased (from yellow to red) and decreased (from light to dark blue) activity for the contrast between emotional reliving of guilt vs. neutral memories.

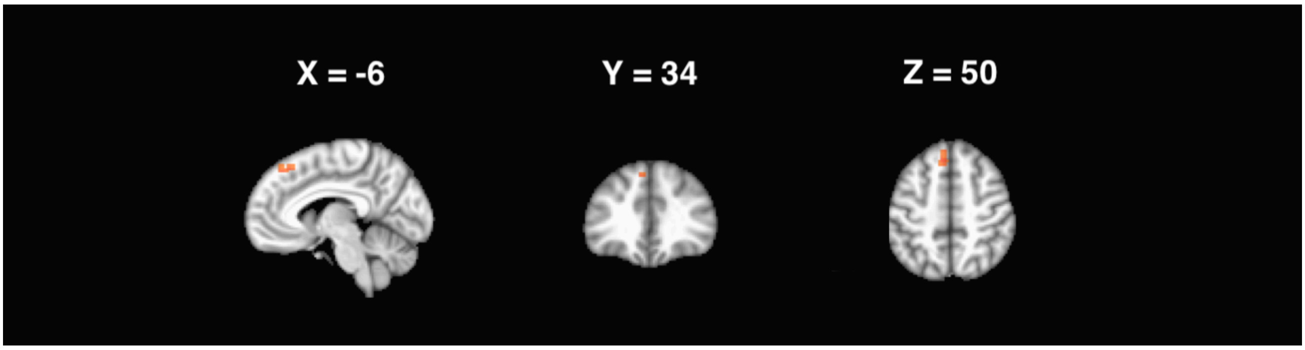


**Supplementary Figure 6.** The left superior frontal gyrus area showed increased task-related functional connectivity with the right superior anterior temporal lobe during emotional reliving of guilt vs. neutral memories.

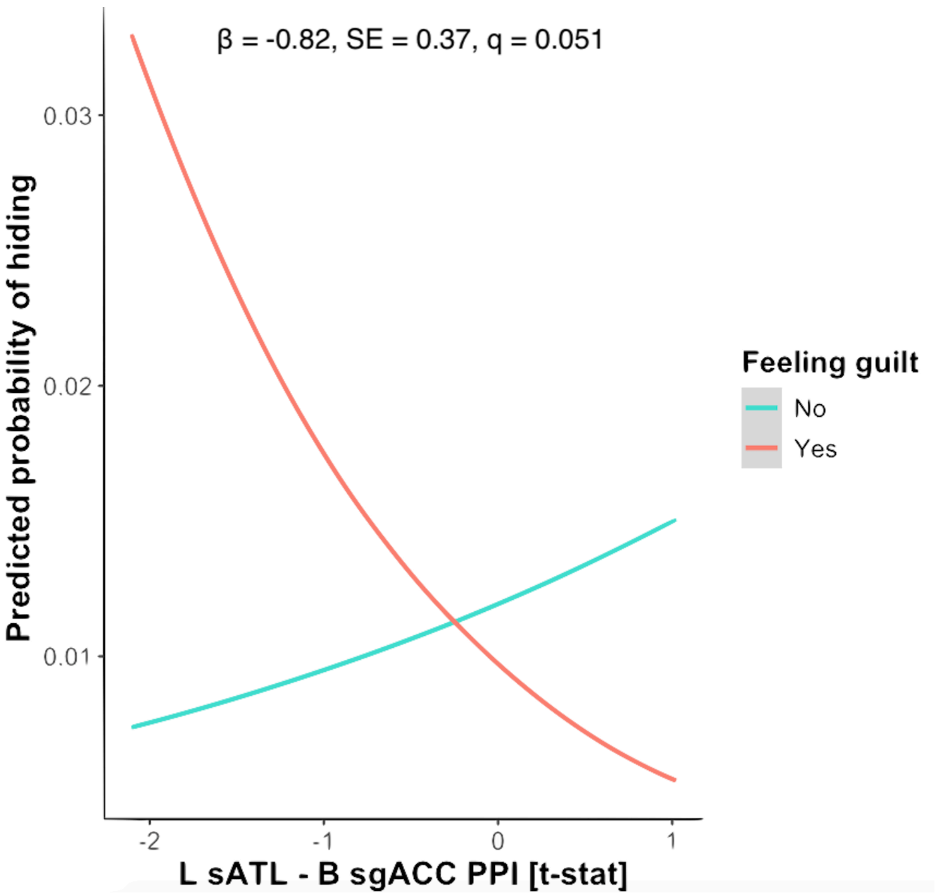


**Supplementary Figure 7.** Individuals with increased task-dependent functional connectivity (PPI) between the left superior anterior temporal lobe (L sATL) and bilateral subgenual anterior cingulate cortex (B sgACC) are less likely to hide when experiencing the feelings of guilt (trend-level). Abbreviations: SE, standard error; q, FDR-corrected p-value.

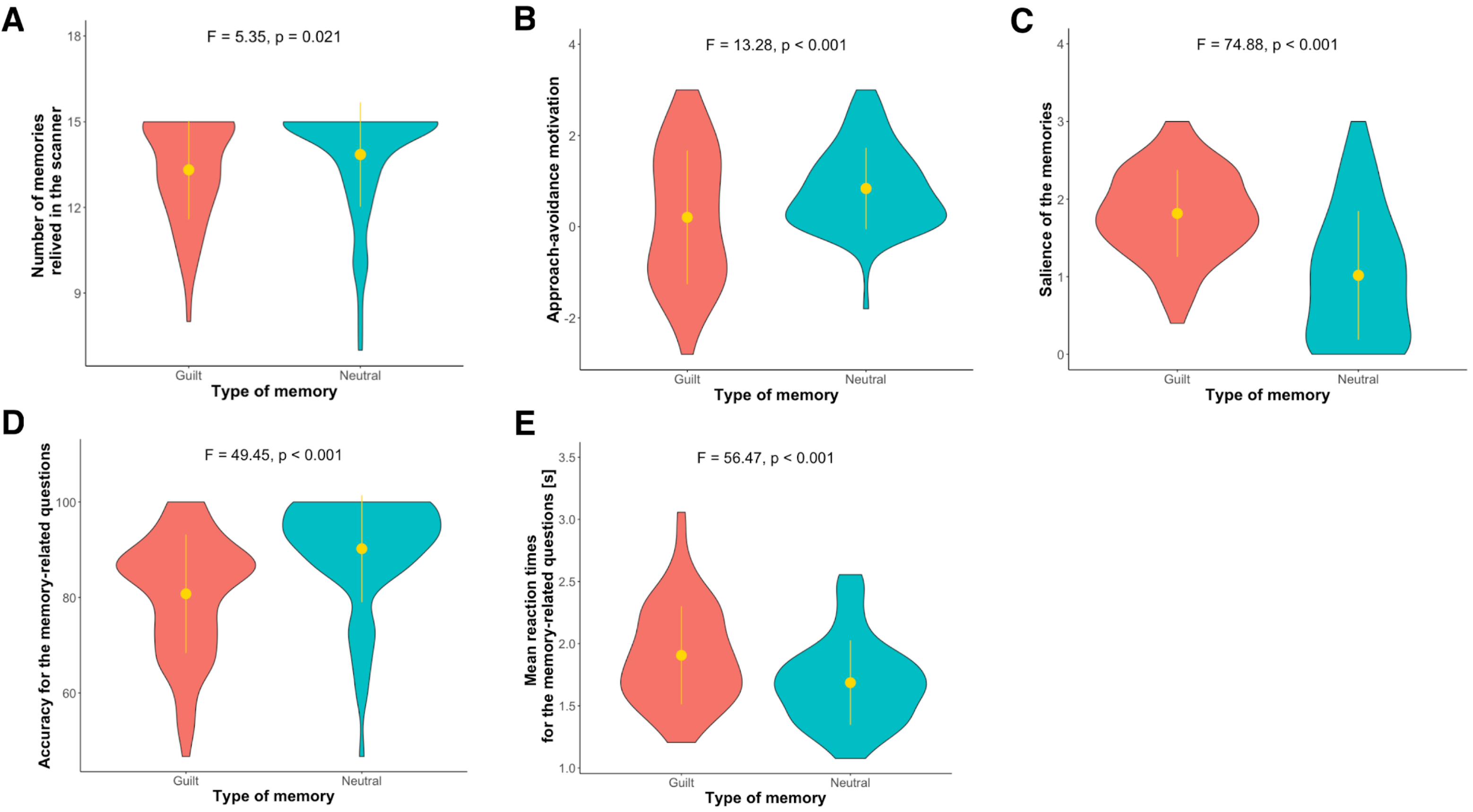


**Supplementary Figure 8.** Measures associated with the guilt recollection task with significant differences between the guilt and neutral memories. Participants found it easier to relive inside the scanner the emotions associated with neutral memories (A). The guilt memories were in turn characterised by less approach/more avoidant motivation (B) and higher salience (C). In the part of the guilt recollection task where the subjects were asked questions regarding the whereabouts of the recalled events, their answers following the neutral memories were more accurate (D) and faster (E). The dots and lines represent, respectively, the means and standard deviations.

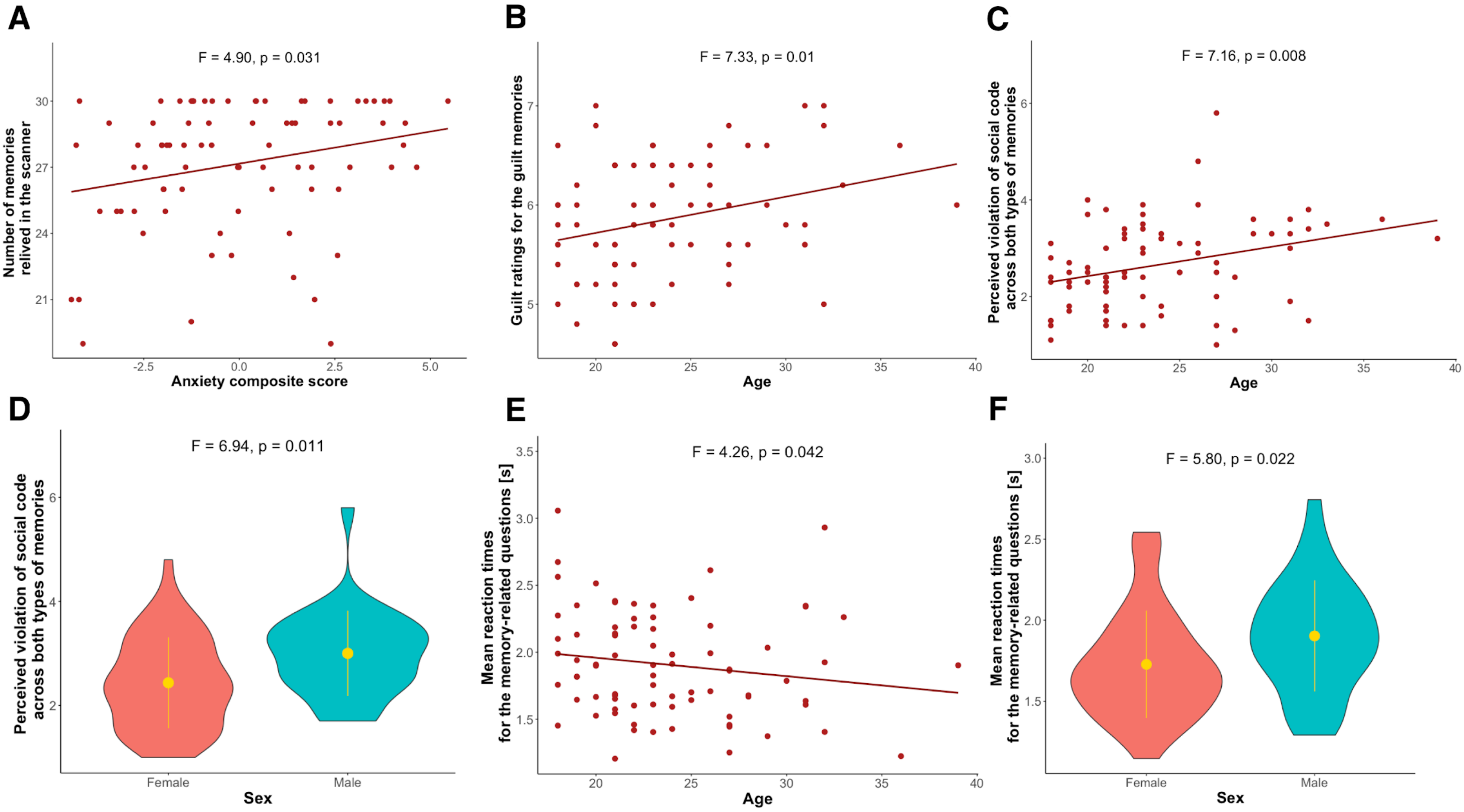


**Supplementary Figure 9.** The associations of anxiety (A), age (B, C, E) and sex (D, F) with the measures associated with the guilt recollection task. Anxiety was positively related to the number of times that participants could successfully relive emotions inside the scanner (A). Older age was in turn related to higher emotional load of the guilt memories (B), stronger perception of themselves violating the social code (C) and lower reaction times when responding to the questions probing whereabouts of their memories (E). The last two characteristics additionally had higher values in males compared to females (D, F). The dots and lines in Figures D and F represent, respectively, the means and standard deviations.

### Exploratory neurochemical correlational analysis

#### 1. Rationale

Despite the fact that both major depressive disorder (MDD) and anxiety disorders patients are often prescribed pharmacotherapies targeting neuromodulatory brain systems, such as serotonin (5-HT), dopamine (DA) and norepinephrine (NE), the neural mechanisms through which these drugs may affect self-blame-related symptomatology remain elusive (Kanen et al., 2021). With the anticipated shift in the paradigm towards personalised psychiatric treatments (Kas et al., 2025), understanding whether particular symptomatology has universal, transdiagnostic underpinnings is essential as such knowledge might help determine the most optimal therapeutic choices. As such, we aimed to investigate whether the guilt processing-evoked brain activity in our sample was related to normative neurochemical densities derived from publicly available atlases (Hansen et al., 2022; Gryglewski et al., 2018). Apart from the aforementioned typical pharmacological targets, we additionally explored the potential role of oxytocin (OXT), given the evidence for its crucial contributions to guilt processing (Li and Zu, 2024; Zheng et al., 2024).

While the gold standard for testing such associations would be collecting both types of data in the same set of participants, its acquisition, for example using positron emission tomography (PET), is costly and requires advanced infrastructure, often unavailable to research centers. The use of normative neurochemical maps, however, opens up the possibility of making preliminary inferences on such associations by performing cross-modality correlations of group-averaged voxel-level values.

#### 2. Methods

##### 2.1. Confirming involvement of neuromodulatory regions in guilt processing

To confirm that the NE, 5-HT and DA systems were involved in guilt processing in our sample, we used publicly available atlases to delineate bilateral locus coeruleus (LC), dorsal raphe (DR) and ventral tegmental area (VTA) in the MNI space (Betts et al., 2017; Beliveau et al., 2015; Murty et al., 2014; see Supplementary Figure E1). The process is described in more detail elsewhere (Zareba et al., 2024a). The quality of the dataset was assessed by obtaining the mean temporal signal-to-noise ratio (tSNR) in each region-of-interest (ROI). The analysis confirmed proper signal coverage for all three areas (LC: 51.78 ± 6.51; DR: 50.33 ± 9.48; VTA: 47.87 ± 6.27), warranting the procedures described below. The mean tSNR map in the entire sample is available at the NeuroVault repository referenced in the main text of the manuscript.

To assess activity of each area during guilt processing, we extracted in each participant the ROI-averaged brain activity for the contrast of reliving of guilt vs. neutral memories, and calculated false discovery rate (FDR)-corrected one sample t-tests. The analysis indicated increased activity of all three neuromodulatory brain regions during guilt processing (Supplementary Figure E2). This confirmatory analysis was not performed for the oxytocin system as the hypothalamic oxytocinergic nuclei could not be properly delineated using the current imaging sequence.

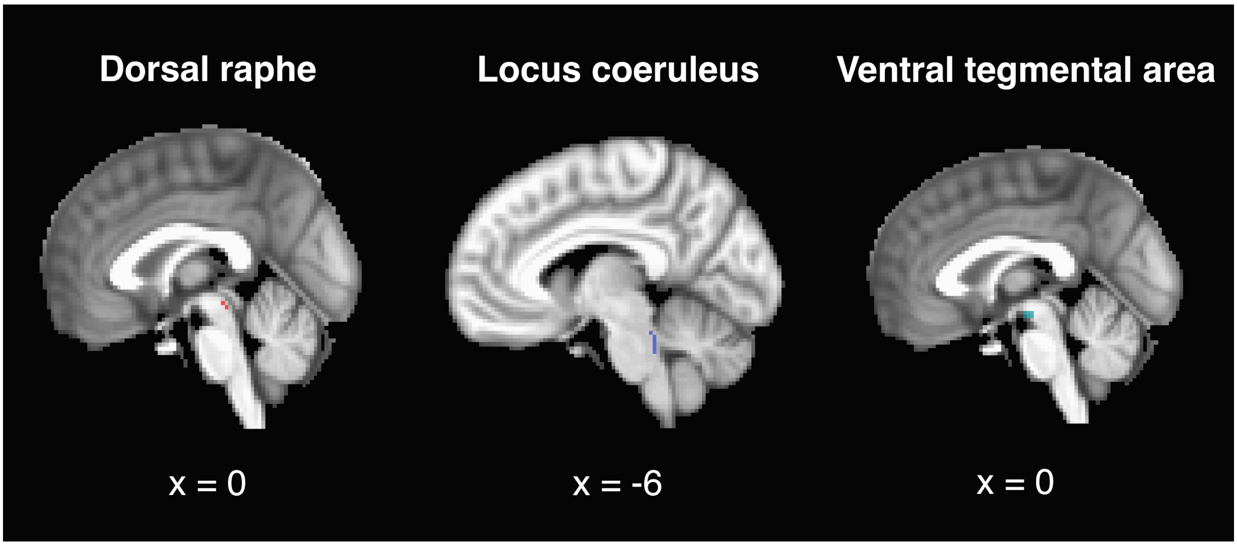


**Supplementary Figure E1.** Delineation of dorsal raphe, locus coeruleus and ventral tegmental area in the MNI space. For visualisation purposes, only the left locus coeruleus was shown.

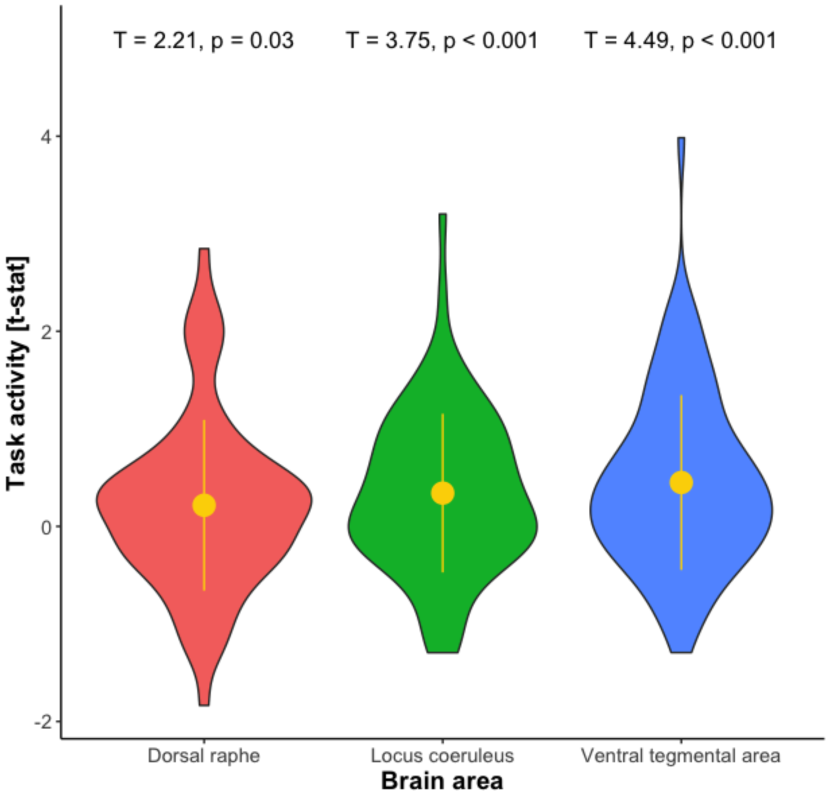


**Supplementary Figure E2.** Results of the one sample t-test performed on the activity of dorsal raphe, bilateral locus coeruleus and ventral tegmental area for the reliving of guilt vs. neutral memories. The tests indicated increased activity for all three areas. The results were corrected for multiple comparisons with the false discovery rate (FDR) method.

### 2.2. Parcellation of brain areas involved in the guilt recollection task

To assess potential interregional variability in their associations with neurochemical densities, we first created a binary mask of areas with significantly higher and lower brain activity associated with guilt processing on the group level, and subsequently used the atlases of cortical, subcortical and cerebellar regions (Fan et al., 2016; Diedrichsen et al., 2009) for its parcellation. As a means of striking a balance between spatial resolution and power to detect meaningful effects within a region, we ensured that each of the parcels contained at least 40 voxels. In the cases when original atlas-based regions contained a smaller number of voxels surviving the cluster-level FWE thresholding, we merged such areas into larger clusters based on the visual convergence of their connectivity patterns and associated cognitive functions, as provided in the atlas (Fan et al., 2016; see Supplementary Table E1 and *Data availability* section in the main manuscript file). For instance, all the unilateral subdivisions of the postcentral gyrus were put together to form one region.

**Supplementary Table E1.** Parcellation of the brain areas with significantly increased or decreased activity for reliving guilt vs. neutral memories.

| Region code | Original parcellation scheme | Original naming of the parcels | Voxels |
|---|---|---|---|
| 1 | Diedrichsen et al., 2009 | B cerebellar lobule I-V | 127 |
| 2 | Diedrichsen et al., 2009 | B cerebellar lobule IX | 73 |
| 3 | Diedrichsen et al., 2009 | B cerebellar lobule VIII | 68 |
| 4 | Diedrichsen et al., 2009 | L cerebellar lobule VI | 140 |
| 5 | Diedrichsen et al., 2009 | L cerebellar crus I | 146 |
| 6 | Diedrichsen et al., 2009 | L cerebellar crus II | 50 |
| 7 | Diedrichsen et al., 2009 | R cerebellar lobule VI | 274 |
| 8 | Diedrichsen et al., 2009 | R cerebellar lobule VIIb | 61 |
| 9 | Diedrichsen et al., 2009 | R cerebellar crus I | 782 |
| 10 | Diedrichsen et al., 2009 | R cerebellar crus II | 378 |
| 11 | Remaining voxels | Pons | 121 |
| 12 | Fan et al., 2016 | L superior frontal gyrus A8m | 244 |
| 13 | Fan et al., 2016 | R superior frontal gyrus A8m | 114 |
| 14 | Fan et al., 2016 | L superior frontal gyrus A8dl | 51 |
| 15 | Fan et al., 2016 | R superior frontal gyrus A8dl<br>R superior frontal gyrus A9l | 185 |
| 16 | Fan et al., 2016 | L superior frontal gyrus A9l | 226 |
| 17 | Fan et al., 2016 | L superior frontal gyrus A6dl | 64 |
| 18 | Fan et al., 2016 | R superior frontal gyrus A6dl | 109 |
| 19 | Fan et al., 2016 | L superior frontal gyrus A6m | 116 |
| 20 | Fan et al., 2016 | R superior frontal gyrus A6m | 83 |
| 21 | Fan et al., 2016 | L superior frontal gyrus A9m | 155 |
| 22 | Fan et al., 2016 | R superior frontal gyrus A9m | 59 |

| | | | |
|---|---|---|---|
| 23 | Fan et al., 2016 | L superior frontal gyrus A10m | 447 |
| 24 | Fan et al., 2016 | R superior frontal gyrus A10m | 236 |
| 25 | Fan et al., 2016 | L middle frontal gyrus A9/46d<br>L middle frontal gyrus A9/46v<br>L middle frontal gyrus A6vl | 41 |
| 26 | Fan et al., 2016 | L middle frontal gyrus A46<br>L middle frontal gyrus IFJ<br>L middle frontal gyrus A10l | 148 |
| 27 | Fan et al., 2016 | R middle frontal gyrus A9/46d | 152 |
| 28 | Fan et al., 2016 | R middle frontal gyrus IFJ | 77 |
| 29 | Fan et al., 2016 | R middle frontal gyrus A46 | 126 |
| 30 | Fan et al., 2016 | R middle frontal gyrus A9/46v | 362 |
| 31 | Fan et al., 2016 | R middle frontal gyrus A8vl | 286 |
| 32 | Fan et al., 2016 | R middle frontal gyrus A6vl | 141 |
| 33 | Fan et al., 2016 | R middle frontal gyrus A10l | 165 |
| 34 | Fan et al., 2016 | L inferior frontal gyrus IFS<br>L inferior frontal gyrus A45c | 111 |
| 35 | Fan et al., 2016 | R inferior frontal gyrus A44d<br>R inferior frontal gyrus IFS | 130 |
| 36 | Fan et al., 2016 | L inferior frontal gyrus A45r | 101 |
| 37 | Fan et al., 2016 | R inferior frontal gyrus A45 | 89 |
| 38 | Fan et al., 2016 | L inferior frontal gyrus A44op | 169 |
| 39 | Fan et al., 2016 | R inferior frontal gyrus A44op<br>R inferior frontal gyrus A44v | 62 |
| 40 | Fan et al., 2016 | L inferior frontal gyrus A44v | 89 |
| 41 | Fan et al., 2016 | L orbital gyrus A14m | 170 |
| 42 | Fan et al., 2016 | R orbital gyrus A14m | 46 |
| 43 | Fan et al., 2016 | L orbital gyrus A12/47o | 77 |
| 44 | Fan et al., 2016 | R orbital gyrus A11l<br>R orbital gyrus A12/47o | 135 |
| 45 | Fan et al., 2016 | L orbital gyrus A13<br>L orbital gyrus A11m | 115 |
| 46 | Fan et al., 2016 | R orbital gyrus A11m | 58 |
| 47 | Fan et al., 2016 | L orbital gyrus A12/47l | 143 |
| 48 | Fan et al., 2016 | R orbital gyrus A12/47l | 135 |

| | | | |
|---|---|---|---|
| 49 | Fan et al., 2016 | L precentral gyrus A4hf<br>L precentral gyrus A6cdl<br>L precentral gyrus A4tl<br>L precentral gyrus A6cvl | 49 |
| 50 | Fan et al., 2016 | R precentral gyrus A6cdl<br>R precentral gyrus A6cvl | 56 |
| 51 | Fan et al., 2016 | L paracentral lobule A1/2/3ll<br>L paracentral lobule A4ll | 135 |
| 52 | Fan et al., 2016 | R paracentral lobule A1/2/3ll<br>R paracentral lobule A4ll | 177 |
| 53 | Fan et al., 2016 | L superior temporal gyrus A38m<br>L superior temporal gyrus A38l | 153 |
| 54 | Fan et al., 2016 | R superior temporal gyrus A38m<br>R superior temporal gyrus A38l | 97 |
| 55 | Fan et al., 2016 | L superior temporal gyrus A22r | 88 |
| 56 | Fan et al., 2016 | L superior temporal gyrus A41/42<br>L superior temporal gyrus TE1.0_TE1.2<br>L superior temporal gyrus A22c | 79 |
| 57 | Fan et al., 2016 | R superior temporal gyrus A41/42<br>R superior temporal gyrus TE1.0_TE1.2<br>R superior temporal gyrus A22c<br>R superior temporal gyrus A22r | 113 |
| 58 | Fan et al., 2016 | L rostroposterior superior temporal sulcus<br>L middle temporal gyrus A37dl<br>L caudoposterior superior temporal sulcus | 112 |
| 59 | Fan et al., 2016 | R rostroposterior superior temporal sulcus | 68 |
| 60 | Fan et al., 2016 | L middle temporal gyrus A21r | 213 |
| 61 | Fan et al., 2016 | L middle temporal gyrus anterior superior temporal sulcus | 293 |
| 62 | Fan et al., 2016 | R middle temporal gyrus A21c | 171 |
| 63 | Fan et al., 2016 | R middle temporal gyrus A21r | 106 |
| 64 | Fan et al., 2016 | R middle temporal gyrus A37dl | 83 |
| 65 | Fan et al., 2016 | R middle temporal gyrus anterior superior temporal sulcus | 94 |
| 66 | Fan et al., 2016 | L inferior temporal gyrus A20iv<br>L inferior temporal gyrus A20r<br>L inferior temporal gyrus A20il<br>L inferior temporal gyrus A20cl<br>L inferior temporal gyrus A20cv<br>L fusiform gyrus A20rv | 172 |
| 67 | Fan et al., 2016 | L inferior temporal gyrus A37elv<br>L inferior temporal gyrus A37vl | 124 |

| | | | |
|---|---|---|---|
| | | L fusiform gyrus A37lv | |
| 68 | Fan et al., 2016 | R inferior temporal gyrus A20r<br>R inferior temporal gyrus A20il<br>R inferior temporal gyrus A20cl<br>R inferior temporal gyrus A20cv<br>R fusiform gyrus A20rv | 242 |
| 69 | Fan et al., 2016 | R inferior temporal gyrus A37elv<br>R inferior temporal gyrus A37vl<br>R fusiform gyrus A37mv | 60 |
| 70 | Fan et al., 2016 | L superior parietal lobule A7r<br>L superior parietal lobule A7c<br>L superior parietal lobule A5l<br>L superior parietal lobule A7pc<br>L superior parietal lobule A7ip | 190 |
| 71 | Fan et al., 2016 | R superior parietal lobule A7c<br>R superior parietal lobule A5l<br>R superior parietal lobule A7ip | 300 |
| 72 | Fan et al., 2016 | L inferior parietal lobule A39c | 155 |
| 73 | Fan et al., 2016 | L inferior parietal lobule A39rd | 64 |
| 74 | Fan et al., 2016 | L inferior parietal lobule A40rd | 255 |
| 75 | Fan et al., 2016 | L inferior parietal lobule A40c | 132 |
| 76 | Fan et al., 2016 | L inferior parietal lobule A39rv | 326 |
| 77 | Fan et al., 2016 | R inferior parietal lobule A40rv<br>L inferior parietal lobule A40rv | 47 |
| 78 | Fan et al., 2016 | R inferior parietal lobule A39c | 64 |
| 79 | Fan et al., 2016 | R inferior parietal lobule A39rv | 56 |
| 80 | Fan et al., 2016 | R inferior parietal lobule A39rd | 414 |
| 81 | Fan et al., 2016 | R inferior parietal lobule A40rd | 418 |
| 82 | Fan et al., 2016 | R inferior parietal lobule A40c | 330 |
| 83 | Fan et al., 2016 | L precuneus A7m<br>L dorsomedial parietooccipital sulcus<br>L precuneus A5m | 104 |
| 84 | Fan et al., 2016 | R precuneus A7m<br>R dorsomedial parietooccipital sulcus | 311 |
| 85 | Fan et al., 2016 | R precuneus A5m | 122 |
| 86 | Fan et al., 2016 | L precuneus A31 | 214 |
| 87 | Fan et al., 2016 | R precuneus A31 | 215 |
| 88 | Fan et al., 2016 | L postcentral gyrus A2<br>L postcentral gyrus A1_2_3_ulhf | 153 |

| | | | |
|---|---|---|---|
| | | L postcentral gyrus A1/2/3tonIa | |
| 89 | Fan et al., 2016 | R postcentral gyrus A1/2/3tonIa<br>R postcentral gyrus A2 | 191 |
| 90 | Fan et al., 2016 | L ventral agranular insular gyrus<br>R ventral agranular insular gyrus<br>L dorsal agranular insular gyrus<br>R dorsal agranular insular gyrus<br>L dorsal dysgranular insular gyrus<br>R dorsal dysgranular insular gyrus | 146 |
| 91 | Fan et al., 2016 | L cingulate gyrus A23d<br>L cingulate gyrus A23v<br>L cingulate gyrus A23c | 137 |
| 92 | Fan et al., 2016 | R cingulate gyrus A23d<br>R cingulate gyrus A23v<br>R cingulate gyrus A23c | 231 |
| 93 | Fan et al., 2016 | L cingulate gyrus A24rv<br>L cingulate gyrus A32sg<br>L cingulate gyrus A32p | 276 |
| 94 | Fan et al., 2016 | R cingulate gyrus A24rv<br>R cingulate gyrus A32p<br>R cingulate gyrus A32sg | 124 |
| 95 | Fan et al., 2016 | L caudal lingual gyrus<br>L caudal cuneus gyrus<br>L lateral occipital cortex V5/MT+<br>L occipital polar cortex<br>L inferior occipital gyrus<br>L lateral superior occipital gyrus<br>L rostral lingual gyrus | 117 |
| 96 | Fan et al., 2016 | R caudal lingual gyrus<br>R ventromedial parietooccipital sulcus<br>R medial superior occipital gyrus<br>R lateral superior occipital gyrus | 171 |
| 97 | Fan et al., 2016 | R parahippocampal gyrus A28_34<br>R parahippocampal gyrus TI_r<br>L medial amygdala<br>R medial amygdala<br>L lateral amygdala<br>R lateral amygdala<br>L rostral hippocampus<br>R rostral hippocampus<br>L caudal hippocampus<br>R caudal hippocampus | 180 |
| 98 | Fan et al., 2016 | L ventral caudate<br>R ventral caudate<br>R globus pallidus<br>L nucleus accumbens<br>R nucleus accumbens<br>L ventromedial putamen<br>L dorsal caudate | 102 |

| | | | |
|---|---|---|---|
| 99 | Fan et al., 2016 | L medial prefrontal thalamus<br>L medial premotor thalamus<br>L rostral temporal thalamus<br>L caudal temporal thalamus | 123 |
| 100 | Fan et al., 2016 | R medial prefrontal thalamus<br>R sensory thalamus<br>R rostral temporal thalamus<br>R posterior parietal thalamus<br>R occipital thalamus<br>R caudal temporal thalamus<br>R lateral prefrontal thalamus | 110 |

Abbreviations: B, bilateral; L, left; R, right.

### 2.3. Neurochemical maps and statistical analysis

With the use of Neuromaps (Hansen et al., 2022), we obtained normative PET maps for:

- dopamine transporter (DAT), as well as $D_1$ and $D_2$ receptors (Dukart et al., 2018; Kaller et al., 2017; Smith et al., 2019; Sandiego et al., 2015; Zakiniaeiz et al., 2019; Slifstein et al., 2015; Sandiego et al., 2019)
- serotonin transporter (5-HTT) as well as $5\text{-HT}_{1B}$, $5\text{-HT}_{2A}$, $5\text{-HT}_4$ and $5\text{-HT}_6$ receptors (Beliveau et al., 2017; Gallezot et al., 2010; Murrough et al., 2011a; Murrough et al., 2011b; Matuskey et al., 2014; Pittenger et al., 2016; Saricicek et al., 2015; Baldassarri et al., 2020; Radhakrishnan et al., 2018; Radhakrishnan et al., 2020; Savli et al., 2012)
- norepinephrine transporter (NET; Ding et al., 2010; Li et al., 2014; Sanchez-Rangel et al., 2020; Belfort-DeAguiar et al., 2018)

As there are no known PET tracers for oxytocin receptors (OXTR), we used the maps of predicted mRNA expression from post-mortem brains (Gryglewski et al., 2018). All the maps of neuromodulatory systems were resampled to the task fMRI data resolution, followed by voxel-wise z-score normalisation. For each ROI, we first extracted for all the voxels (group-level) t-stats from the contrast map of interest and Z-scores from the external maps of neuromodulatory molecules densities, and subsequently used Spearman correlations to probe the cross-modality associations. FDR (< 0.05) was used to correct for the number of regions (N = 100), separately for each molecular map. Of note, the current analysis did not account for the spatial autocorrelation of the neurochemical maps. The results presented below should be therefore treated as purely descriptive.

### 3. Results

The analyses revealed robust associations for all the tested neuromodulatory systems, i.e. 5-HT, DA, NE and OXT, indicating that the discussed neurotransmitters might play a crucial role in shaping the neural dynamics when one experiences strong and negative self-referential emotions (see Figure 3 in the main text of the manuscript). Thresholded correlation maps for specific molecules are available on the NeuroVault platform (see *Data availability* section in the main text of the manuscript).

Interestingly, the results hint at a possibility of differential involvement of specific neurotransmitters across the brain, even if the areas form part of a tightly cooperating network. For instance, the activity in the right-sided dorsolateral prefrontal cortex and anterior inferior parietal lobule, mapping onto the lateral frontoparietal control network, crucial for voluntary emotion regulation (Morawetz et al., 2020), was correlated with indices of all 4 neuromodulatory systems, while the link with NE was absent for the more posterior parts of the inferior parietal lobule. Similarly, the activity in the bilateral hippocampi and amygdalae, key regions for emotional memory, was associated solely with 5-HT and OXT molecules density, while the insula, another region strongly implicated in emotion processing and etiology of mood and anxiety disorders (Picó-Pérez et al., 2017; Zareba et al., 2024b; Zareba et al., 2025), displayed a more complex, subregion-specific pattern.

Gryglewski G, Seiger R, James GM, Godbersen GM, Komorowski A, Unterholzner J, Michenthaler P, Hahn A, Wadsak W, Mitterhauser M, Kasper S, Lanzenberger R. Spatial analysis and high resolution mapping of the human whole-brain transcriptome for integrative analysis in neuroimaging.

Hansen JY, Shafiei G, Markello RD, Smart K, Cox SML, Nørgaard M, Beliveau V, Wu Y, Gallezot JD, Aumont É, Servaes S, Scala SG, DuBois JM, Wainstein G, Bezgin G, Funck T, Schmitz TW, Spreng RN, Galovic M, Koepp MJ, Duncan JS, Coles JP, Fryer TD, Aigbirhio FI, McGinnity CJ, Hammers A, Soucy JP, Baillet S, Guimond S, Hietala J, Bedard MA, Leyton M, Kobayashi E, Rosa-Neto P, Ganz M, Knudsen GM, Palomero-Gallagher N, Shine JM, Carson RE, Tuominen L, Dagher A, Misic B. Mapping neurotransmitter systems to the structural and functional

organization of the human neocortex. Nat Neurosci. 2022 Nov;25(11):1569-1581. doi: 10.1038/s41593-022-01186-3.
Kaller S, Rullmann M, Patt M, Becker GA, Luthardt J, Girbardt J, Meyer PM, Werner P, Barthel H, Bresch A, Fritz TH, Hesse S, Sabri O. Test-retest measurements of dopamine D1-type receptors using simultaneous PET/MRI imaging. Eur J Nucl Med Mol Imaging. 2017 Jun;44(6):1025-1032. doi: 10.1007/s00259-017-3645-0.
Kanen JW, Arntz FE, Yellowlees R, Cardinal RN, Price A, Christmas DM, Apergis-Schoute AM, Sahakian BJ, Robbins TW. Serotonin depletion amplifies distinct human social emotions as a function of individual differences in personality. Transl Psychiatry. 2021 Feb 1;11(1):81. doi: 10.1038/s41398-020-00880-9.
Kas MJH, Penninx BWJH, Knudsen GM, Cuthbert B, Falkai P, Sachs GS, Ressler KJ, Bałkowiec-Iskra E, Butlen-Ducuing F, Leboyer M, Marston H, Luthman J, Mantua V. Precision psychiatry roadmap: towards a biology-informed framework for mental disorders. Mol Psychiatry. 2025 Aug;30(8):3846-3855. doi: 10.1038/s41380-025-03070-5.
Li CS, Potenza MN, Lee DE, Planeta B, Gallezot JD, Labaree D, Henry S, Nabulsi N, Sinha R, Ding YS, Carson RE, Neumeister A. Decreased norepinephrine transporter availability in obesity: Positron Emission Tomography imaging with (S,S)-[(11)C]O-methylreboxetine. Neuroimage. 2014 Feb 1;86:306-10. doi: 10.1016/j.neuroimage.2013.10.004.
Li Z, Xu M. Oxytocin enhances group-based guilt in high moral disengagement individuals through increased moral responsibility. Psychoneuroendocrinology. 2024 Oct;168:107131. doi: 10.1016/j.psyneuen.2024.107131.
Matuskey D, Bhagwagar Z, Planeta B, Pittman B, Gallezot JD, Chen J, Wanyiri J, Najafzadeh S, Ropchan J, Geha P, Huang Y, Potenza MN, Neumeister A, Carson RE, Malison RT. Reductions in brain 5-HT1B receptor availability in primarily cocaine-dependent humans. Biol Psychiatry. 2014 Nov 15;76(10):816-22. doi: 10.1016/j.biopsych.2013.11.022.
Morawetz C, Riedel MC, Salo T, Berboth S, Eickhoff SB, Laird AR, Kohn N. Multiple large-scale neural networks underlying emotion regulation. Neurosci Biobehav Rev. 2020 Sep;116:382-395. doi: 10.1016/j.neubiorev.2020.07.001.
Murrough JW, Czermak C, Henry S, Nabulsi N, Gallezot JD, Gueorguieva R, Planeta-Wilson B, Krystal JH, Neumaier JF, Huang Y, Ding YS, Carson RE, Neumeister A. The effect of early trauma exposure on serotonin type 1B receptor expression revealed by reduced selective radioligand binding. Arch Gen Psychiatry. 2011a Sep;68(9):892-900. doi: 10.1001/archgenpsychiatry.2011.91.
Murrough JW, Henry S, Hu J, Gallezot JD, Planeta-Wilson B, Neumaier JF, Neumeister A. Reduced ventral striatal/ventral pallidal serotonin1B receptor binding potential in major depressive disorder. Psychopharmacology (Berl). 2011b Feb;213(2-3):547-53. doi: 10.1007/s00213-010-1881-0.
Murty VP, Shermohammed M, Smith DV, Carter RM, Huettel SA, Adcock RA. Resting state networks distinguish human ventral tegmental area from substantia nigra. Neuroimage. 2014 Oct 15;100:580-9. doi: 10.1016/j.neuroimage.2014.06.047.
Picó-Pérez M, Radua J, Steward T, Menchón JM, Soriano-Mas C. Emotion regulation in mood and anxiety disorders: A meta-analysis of fMRI cognitive reappraisal studies. Prog Neuropsychopharmacol Biol Psychiatry. 2017 Oct 3;79(Pt B):96-104. doi: 10.1016/j.pnpbp.2017.06.001.
Pittenger C, Adams TG Jr, Gallezot JD, Crowley MJ, Nabulsi N, James Ropchan, Gao H, Kichuk SA, Simpson R, Billingslea E, Hannestad J, Bloch M, Mayes L, Bhagwagar Z, Carson RE. OCD is associated with an altered association between sensorimotor gating and cortical and subcortical 5-HT1b receptor binding. J Affect Disord. 2016 May 15;196:87-96. doi: 10.1016/j.jad.2016.02.021.
Radhakrishnan R, Matuskey D, Nabulsi N, Gaiser E, Gallezot JD, Henry S, Planeta B, Lin SF, Ropchan J, Huang Y, Carson RE, D'Souza DC. In vivo 5-HT6 and 5-HT2A receptor availability in antipsychotic treated schizophrenia patients vs. unmedicated healthy humans measured with

[11C]GSK215083 PET. Psychiatry Res Neuroimaging. 2020 Jan 30;295:111007. doi: 10.1016/j.pscychresns.2019.111007.
Radhakrishnan R, Nabulsi N, Gaiser E, Gallezot JD, Henry S, Planeta B, Lin SF, Ropchan J, Williams W, Morris E, D'Souza DC, Huang Y, Carson RE, Matuskey D. Age-Related Change in 5-HT6 Receptor Availability in Healthy Male Volunteers Measured with 11C-GSK215083 PET. J Nucl Med. 2018 Sep;59(9):1445-1450. doi: 10.2967/jnumed.117.206516.
Sanchez-Rangel E, Gallezot JD, Yeckel CW, Lam W, Belfort-DeAguiar R, Chen MK, Carson RE, Sherwin R, Hwang JJ. Norepinephrine transporter availability in brown fat is reduced in obesity: a human PET study with [11C] MRB. Int J Obes (Lond). 2020 Apr;44(4):964-967. doi: 10.1038/s41366-019-0471-4.
Sandiego CM, Gallezot JD, Lim K, Ropchan J, Lin SF, Gao H, Morris ED, Cosgrove KP. Reference region modeling approaches for amphetamine challenge studies with [11C]FLB 457 and PET. J Cereb Blood Flow Metab. 2015 Mar 31;35(4):623-9. doi: 10.1038/jcbfm.2014.237.
Sandiego CM, Matuskey D, Lavery M, McGovern E, Huang Y, Nabulsi N, Ropchan J, Picciotto MR, Morris ED, McKee SA, Cosgrove KP. The Effect of Treatment with Guanfacine, an Alpha2 Adrenergic Agonist, on Dopaminergic Tone in Tobacco Smokers: An [11C]FLB457 PET Study. Neuropsychopharmacology. 2018 Apr;43(5):1052-1058. doi: 10.1038/npp.2017.223.
Saricicek A, Chen J, Planeta B, Ruf B, Subramanyam K, Maloney K, Matuskey D, Labaree D, Deserno L, Neumeister A, Krystal JH, Gallezot JD, Huang Y, Carson RE, Bhagwagar Z. Test-retest reliability of the novel 5-HT1B receptor PET radioligand [11C]P943. Eur J Nucl Med Mol Imaging. 2015 Mar;42(3):468-77. doi: 10.1007/s00259-014-2958-5.
Savli M, Bauer A, Mitterhauser M, Ding YS, Hahn A, Kroll T, Neumeister A, Haeusler D, Ungersboeck J, Henry S, Isfahani SA, Rattay F, Wadsak W, Kasper S, Lanzenberger R. Normative database of the serotonergic system in healthy subjects using multi-tracer PET. Neuroimage. 2012 Oct 15;63(1):447-59. doi: 10.1016/j.neuroimage.2012.07.001.
Slifstein M, van de Giessen E, Van Snellenberg J, Thompson JL, Narendran R, Gil R, Hackett E, Girgis R, Ojeil N, Moore H, D'Souza D, Malison RT, Huang Y, Lim K, Nabulsi N, Carson RE, Lieberman JA, Abi-Dargham A. Deficits in prefrontal cortical and extrastriatal dopamine release in schizophrenia: a positron emission tomographic functional magnetic resonance imaging study. JAMA Psychiatry. 2015 Apr;72(4):316-24. doi: 10.1001/jamapsychiatry.2014.2414.
Smith CT, Crawford JL, Dang LC, Seaman KL, San Juan MD, Vijay A, Katz DT, Matuskey D, Cowan RL, Morris ED, Zald DH, Samanez-Larkin GR. Partial-volume correction increases estimated dopamine D2-like receptor binding potential and reduces adult age differences. J Cereb Blood Flow Metab. 2019 May;39(5):822-833. doi: 10.1177/0271678X17737693.
Zakiniaeiz Y, Hillmer AT, Matuskey D, Nabulsi N, Ropchan J, Mazure CM, Picciotto MR, Huang Y, McKee SA, Morris ED, Cosgrove KP. Sex differences in amphetamine-induced dopamine release in the dorsolateral prefrontal cortex of tobacco smokers. Neuropsychopharmacology. 2019 Dec;44(13):2205-2211. doi: 10.1038/s41386-019-0456-y.
Zareba MR, Ariño-Braña P, Picó-Pérez M, Visser M. Noradrenergic and Dopaminergic Neural Correlates of Trait Anxiety: Unveiling the Impact of Maladaptive Emotion Regulation. bioRxiv 2024a 2024.07.23.604801, doi: 10.1101/2024.07.23.604801.
Zareba MR, Bielski K, Costumero V, Visser M. Graph analysis of guilt processing network highlights links with subclinical anxiety and self-blame. Soc Cogn Affect Neurosci. 2024b Dec 13;19(1):nsae092. doi: 10.1093/scan/nsae092.
Zareba MR, Davydova T, Palomar-García MÁ, Adrián-Ventura J, Costumero V, Visser M. Subjective sleep quality in healthy young adults moderates associations of sensitivity to punishment and reward with functional connectivity of regions relevant for insomnia disorder. Sleep Med. 2025 Jul;131:106527. doi: 10.1016/j.sleep.2025.106527.
Zheng X, Wang J, Yang X, Xu L, Becker B, Sahakian BJ, Robbins TW, Kendrick KM. Oxytocin, but not vasopressin, decreases willingness to harm others by promoting moral emotions of guilt and shame. Mol Psychiatry. 2024 Nov;29(11):3475-3482. doi: 10.1038/s41380-024-02590-w
